# A-type FHFs mediate resurgent currents through TTX-resistant voltage-gated sodium channels

**DOI:** 10.1101/2022.03.04.482974

**Authors:** Yucheng Xiao, Jonathan W. Theile, Agnes Zybura, Yanling Pan, Zhixin Lin, Theodore R. Cummins

## Abstract

Resurgent currents (*I*_NaR_) produced by voltage-gated sodium channels are required for many neurons to maintain high-frequency firing, and contribute to neuronal hyperexcitability and disease pathophysiology. Here we show, for the first time, that *I*_NaR_ can be reconstituted in a heterologous system by co-expression of sodium channel α-subunits and A-type fibroblast growth factor homologous factors (FHFs). Specifically, A-type FHFs induces *I*_NaR_ from Nav1.8, Nav1.9 tetrodotoxin-resistant neuronal channels and, to a lesser extent, neuronal Nav1.7 and cardiac Nav1.5 channels. Moreover, we identified the N-terminus of FHF as the critical molecule responsible for A-type FHFs-mediated *I*_NaR_. Among the FHFs, FHF4A is the most important isoform for mediating Nav1.8 and Nav1.9 *I*_NaR_. In nociceptive sensory neurons, FHF4A knockdown significantly reduces *I*_NaR_ amplitude and the percentage of neurons that generate *I*_NaR_, substantially suppressing excitability. Thus, our work reveals a novel molecular mechanism underlying TTX-resistant *I*_NaR_ generation and provides important potential targets for pain treatment.

## Introduction

Voltage-gated sodium channels (VGSCs) are crucial determinants of action potentials in almost all excitable tissues. VGSCs are composed of a functional pore-forming α-subunit associated with auxiliary β-subunits (Catterall et al., 2005). VGSCs also interact with other intracellular proteins, such as fibroblast growth factor homologous factors (FHFs) and calmodulin (Catterall et al., 2005; Wildburger et al., 2015). Although the α-subunit is sufficient to produce a functional VGSC, interacting partners can influence multiple properties of the α-subunits, regulating neuronal excitability (Namadurai et al., 2015). One of the most striking influences is generation of resurgent sodium currents (*I*_NaR_) (Lewis and Raman, 2014).

*I*_NaR_ were originally observed in cerebellar Purkinje neurons (Raman and Bean, 1997), and have been identified in cerebellum, brainstem, trigeminal ganglia and dorsal root ganglion (DRG) neurons (Afshari et al., 2004, Enomoto et al., 2006; Kim et al., 2010). *I*_NaR_ can enhance high-frequency firing in many neurons (Raman and Bean, 1997, Xie et al., 2016), and aberrant *I*_NaR_ have been implicated in multiple human diseases including pain disorders (Jarecki et el., 2010; Patel et al., 2016; Theile et al., 2011; Tanaka et al., 2016). Unlike classic sodium currents that are activated during the depolarizing phase of action potentials, *I*_NaR_ are atypical sodium currents evoked during the repolarizing phase. Navβ4 has been implicated as a major contributor to *I*_NaR_ generation (Grieco et al., 2005; Barbosa et al., 2015; Cannon and Bean, 2010). The most direct evidence supporting the Navβ4 mechanism is that a short peptide derived from the C-terminal tail of Navβ4 can reconstitute the *I*_NaR_. However, the Navβ4 mechanism remains controversial for at least two reasons: (1) Navβ4 knockout or knockdown does not abolish *I*_NaR_ in central (White et al., 2019; Ransdell et al., 2017) or peripheral neurons (Xiao et al., 2019), but rather results in only partial to no reduction of *I*_NaR_; and (2) importantly, coexpression of full-length Navβ4 with VGSC α-subunits fails to reconstitute *I*_NaR_ in heterologous systems. Therefore, other molecular mechanisms for *I*_NaR_ generation remain to be uncovered.

FHFs are widely distributed throughout the CNS/PNS. They represent an important group of auxiliary VGSC subunits that influence neuronal excitability. FHF is a subfamily of the fibroblast growth factor (FGF) superfamily. They can bind to the VGSC C-terminal tails and can modulate VGSCs functional properties, trafficking and axonal localization (Liu et al., 2001; Goetz et al., 2009; Wittmack et al., 2004; Lou et al., 2005; Wang et al., 2011b). There are two main types of FHFs: A-type and B-type. The former has four isoforms (FHF1A – FHF4A). There is emerging evidence that FHFs regulate *I*_NaR_ generation in neurons. In DRG neurons, overexpression of FHF2A and FHF2B decreases and increases Nav1.6 *I*_NaR_, respectively (Barbosa et al., 2017). In contrast FHF4A, which has high sequence similarity to FHF2A, has been proposed to directly mediate *I*_NaR_ generation by Nav1.6. FHF4 knockout significantly reduced *I*_NaR_ in Purkinje neurons, which is mainly carried by Nav1.6, and a peptide corresponding to FHF4A residues 50-63 induced robust *I*_NaR_ in CA3 neurons (White et al., 2019). However, both FHF2A and FHF4A have been shown to induce accumulation of rapid onset long-term inactivation when coexpressed with Nav1.6 in heterologous systems (Venkatesan et al., 2014; Dover et al., 2010). In addition, the reduction in cerebellar Purkinje neuron *I*_NaR_ with FHF4 knockdown has been proposed to be due to an indirect effect involving FHF4B modulation of channel inactivation (Yan et al., 2014). Therefore, there is a lack of compelling evidence supporting a specific molecular mechanism of *I*_NaR_ generation.

In this study, we report that A-type FHFs directly mediate resurgent sodium current generation in Nav1.8 and Nav1.9 sensory neuron VGSCs, and show for the first time that *I*_NaR_ can be reconstituted in heterologous systems by coexpressing full-length A-type FHFs with VGSC α-subunits. These FHF-mediated *I*_NaR_ are independent of Navβ4. The novel FHF-mediated *I*_NaR_ could be fully reproduced by the amino acids 2-21 from the A-type FHF N-terminus. We also show that while FHF2A could induce small *I*_NaR_ with Nav1.5 and Nav1.7, FHF4A did not induce Nav1.5, Nav1.6 or Nav1.7 *I*_NaR_ in heterologous expression systems. We further show that reduction of FHF4A-mediated TTX-resistant *I*_NaR_ substantially downregulated excitability of nociceptive DRG neurons. Because Nav1.7 - Nav1.9 are predominantly expressed in neurons of DRG and trigeminal ganglia, and are crucial for pain perception and transmission (Cummins et al., 2004; Cox et al., 2006; Huang et al., 2014; Dib-Hajj et al., 2015; Huang et al., 2013; Cummins et al., 2007; Dib-Hajj et al., 2010), our work not only uncovers a novel mechanism of *I*_NaR_ generation in sensory neurons, but also identifies an exciting target for the development of new pain treatments.

## Results

### A-type FHFs mediate *I*_NaR_ in heterologously expressed Nav1.8 and Nav1.9

A-type FHFs can modulate TTX-sensitive VGSC inactivation and *I*_NaR_ (White et al., 2019; Barbosa et al., 2017; Yan et al., 2014), however it is unknown if A-type FHFs impact the functional properties of the TTX-resistant sodium channels Nav1.8 and Nav1.9. Therefore, we first asked whether FHF2A and FHF4A, which are widely expressed in DRG neurons, modulate sodium currents in cells expressing recombinant Nav1.8 and Nav1.9. As previously shown in heterologous systems (Xiao et al., 2019; Lin et al., 2016), Nav1.8 generated a slow-inactivating TTX-resistant current, while Nav1.9 produced an ultra-slow-inactivating TTX-resistant current that activated at hyperpolarized potentials (Fig. 1*a*, *d*). Here we show that FHF2A, FHF2B and FHF4A, when coexpressed with Nav1.8, shifted the voltage-dependence of activation by > 7 mV in the negative direction, and shifted the voltage dependence of steady-state inactivation by > 15 mV in the positive direction (Fig. 1*b*; Table 1). FHF2A, FHF2B and FHF4A also accelerated recovery rate from inactivation of Nav1.8 (Fig. 1*c*). When coexpressed with Nav1.9, FHF2A and FHF4A, but not FHF2B, positively shifted the voltage-dependence of steady-state inactivation by ∼10 mV. Distinct from Nav1.8, none of the three FHF isoforms altered the voltage-dependence of activation or rate for recovery from inactivation of Nav1.9 (Fig. 1*e*, *f*; Table 1), suggesting that FHFs differentially regulate TTX-resistant VGSCs.

**Figure 1.**
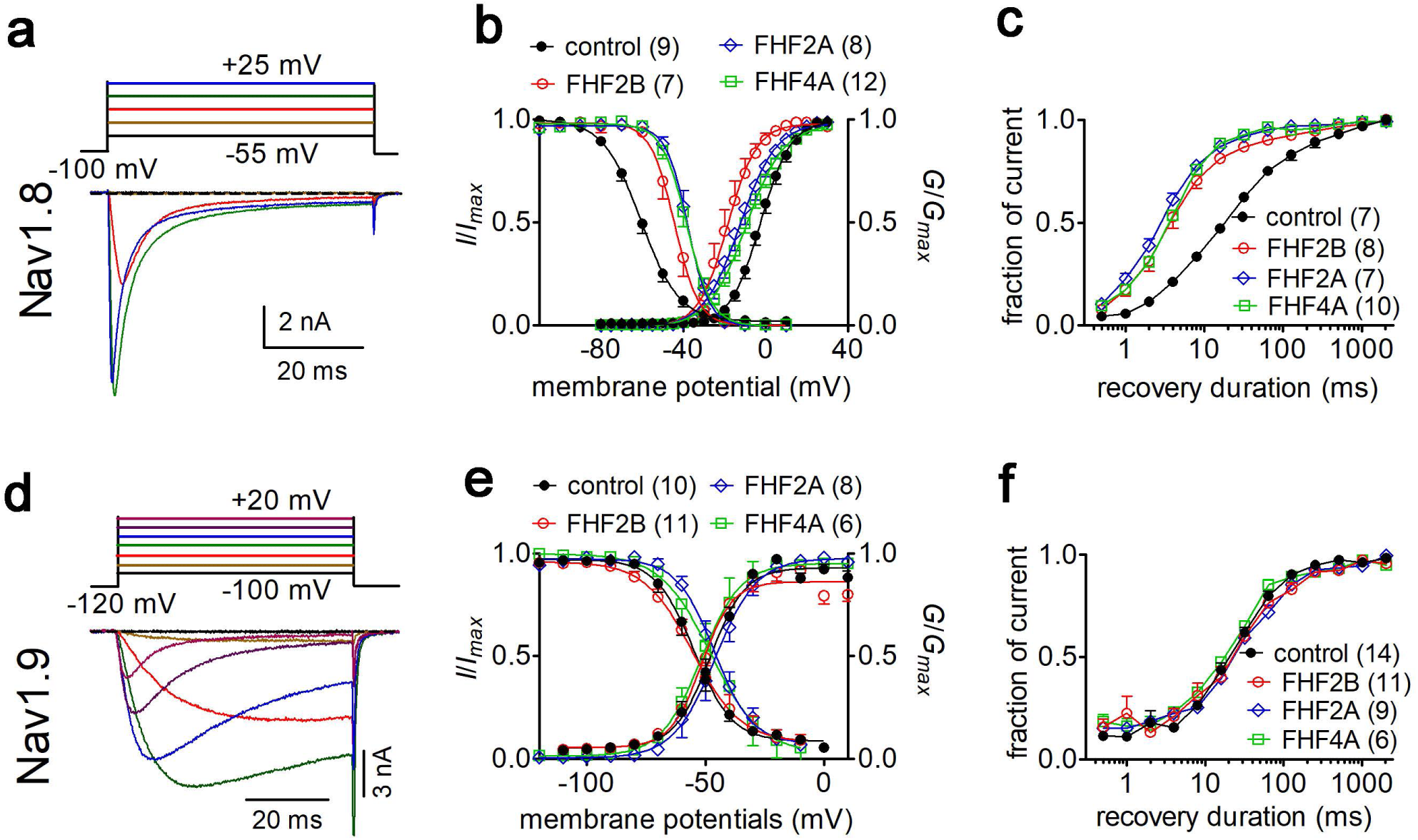
FHFs differentially modulated the gating properties of Nav1.8 and Nav1.9 in heterologous systems. (**a**) Family of classical currents recorded from ND7/23 cells expressing recombinant Nav1.8. Currents were elicited by 50-ms depolarizing voltage steps from +25 mV to −55 mV in -10 mV increments from a holding potential of -100 mV (*inset*). (**b**) Effects of FHF2B, FHF2A and FHF4A on steady-state activation (p < 0.0001, 0.0035, 0.0077 *vs* control, respectively) and inactivation (p = 0.0002, < 0.0001, < 0.0001 *vs* control, respectively) of Nav1.8. (**c**) FHF2B, FHF2A and FHF4A accelerated the recovery rate from Nav1.8 inactivation. The time constants estimated from single exponential fits were 29.71± 2.54 ms (control), 5.81 ± 1.03 ms (FHF2B, p < 0.0001 *vs* control), 4.45 ± 0.43 ms (FHF2A, p < 0.0001 *vs* control) and 5.46 ± 0.40 ms (FHF4A, p < 0.0001 *vs* control). (**d**) Family of classical currents recorded from HEK293 cells expressing recombinant Nav1.9. Currents were elicited by 50-ms depolarizing voltage steps from +20 mV to −100 mV in -20 mV increments from a holding potential of - 120 mV (inset). (**e**) Effects of FHF2B, FHF2A and FHF4A on steady-state activation (p = 0.1832, 0.0171, 0.3215 *vs* control, respectively) and inactivation (p = 0.175, 0.5978, 0.636 *vs* control, respectively) of Nav1.9. (**f**), FHF2B, FHF2A and FHF4A did not affect the recovery rate from Nav1.9 inactivation. The time constants estimated from single exponential fits were 38.46 ± 4.64 ms (control), 48.99 ± 6.93 ms (FHF2B, p = 0.2041 *vs* control), 49.72 ± 6.81 ms (FHF2A, p = 0.1745 *vs* control) and 31.95 ± 2.84 ms (FHF4A, p = 0.4786 *vs* control). In (**a**-**c**), cells were pretreated with 500 nM TTX. Filled circles, open circles, open diamond and open squares represent control, FHF2B, FHF2A and FHF4A, respectively. The number of separate cells tested is indicated in parentheses. Data points are shown as mean ± S.E. The V_1/2_ values for activation and inactivation are summarized in Table 1.

**Table 1.** Gating properties of Nav1.8 and Nav1.9 in the presence of FHFs

| construct | $V_{1/2}$ (mV) | control | FHF2A | FHF2B | FHF4A |
| --- | --- | --- | --- | --- | --- |
| Nav1.8 | activation | $-2.4 \pm 1.5$ (9) | $-12.4 \pm 2.7^{\text{a}}$ (8) | $-18.5 \pm 2.7^{\text{a}}$ (7) | $-9.0 \pm 1.9^*$ (12) |
| | inactivation | $-59.6 \pm 1.8$ (9) | $-37.9 \pm 1.9^{\text{a}}$ (8) | $-44.3 \pm 2.3^{\text{a}}$ (7) | $-39.0 \pm 2.2^{\text{a}}$ (12) |
| Nav1.9 | activation | $-47.8 \pm 2.5$ (10) | $-46.0 \pm 1.2$ (8) | $-50.9 \pm 2.0$ (11) | $-50.5 \pm 4.8$ (6) |
| | inactivation | $-55.2 \pm 2.3$ (10) | $-45.9 \pm 2.5^*$ (8) | $-53.2 \pm 4.2$ (11) | $-45.2 \pm 4.3$ (6) |

We next examined the effects of FHFs on *I*_NaR_ generation. FHF2A and FHF4A induced robust *I*_NaR_ from Nav1.8 and Nav1.9 in HEK293 cells (Fig. 2*a*-*h*). However, under control conditions and with coexpression of FHF2B, the repolarization pulses only elicited classic tail currents, which arise nearly instantaneously and decay rapidly, in Nav1.8 and Nav1.9 (Fig. 2*a* (*left*), *b*, *e* (*left*) and *f*). This is the first demonstration of *I*_NaR_ generation in a heterologous expression system without inclusion of an exogenous peptide in the intracellular solution. The FHF-mediated Nav1.8 *I*_NaR_ peaked at -20 to -10 mV and could be observed at repolarization pulses ranging from +5 to -80 mV, while the FHF-mediated Nav1.9 *I*_NaR_ displayed a more hyperpolarized voltage dependence, peaking at -85 mV and observed at repolarizing potentials ranging from -55 to -100 mV (Fig. 2*e,g*). Moreover, the Nav1.8 *I*_NaR_ induced by FHF4A were four-fold larger than those by FHF2A (Fig. 2*C*, relative amplitudes of the peak transient current: FHF2A: 1.5% ± 0.2%; FHF4A: 5.9% ± 0.4%), and the Nav1.9 *I*_NaR_ mediated by FHF4A was two-fold larger than those induced by FHF2A (FHF2A: 14.8% ± 3.9%; FHF4A: 32.6% ± 3.0%). The kinetics of Nav1.8 *I*_NaR_ mediated by A-type FHF are slow, with a slow onset and slow decay. The time to peak and the decay time constant for the FHF4A-mediated *I*_NaR_ elicited at -20 mV were 9.63 ± 0.61 ms and 85.97 ± 5.29 ms, respectively (Fig. 2*d*), similar to the TTX-resistant *I*_NaR_ previously recorded from DRG neurons (19). In contrast, FHF-mediated Nav1.9 *I*_NaR_ exhibit fast onset and decay kinetics. At -70 mV, near the physiological resting membrane potential of DRG neurons, the time to peak and the decay time constant for FHF4A-mediated Nav1.9 *I*_NaR_ were 1.92 ± 0.12 ms and 8.09 ± 1.31 ms, respectively (Fig. 2*h*). It is noteworthy that Nav1.9 produced a non-decaying inward current following the *I*_NaR_ (control, Fig. 2*e*-*f*).

**Figure 2.**
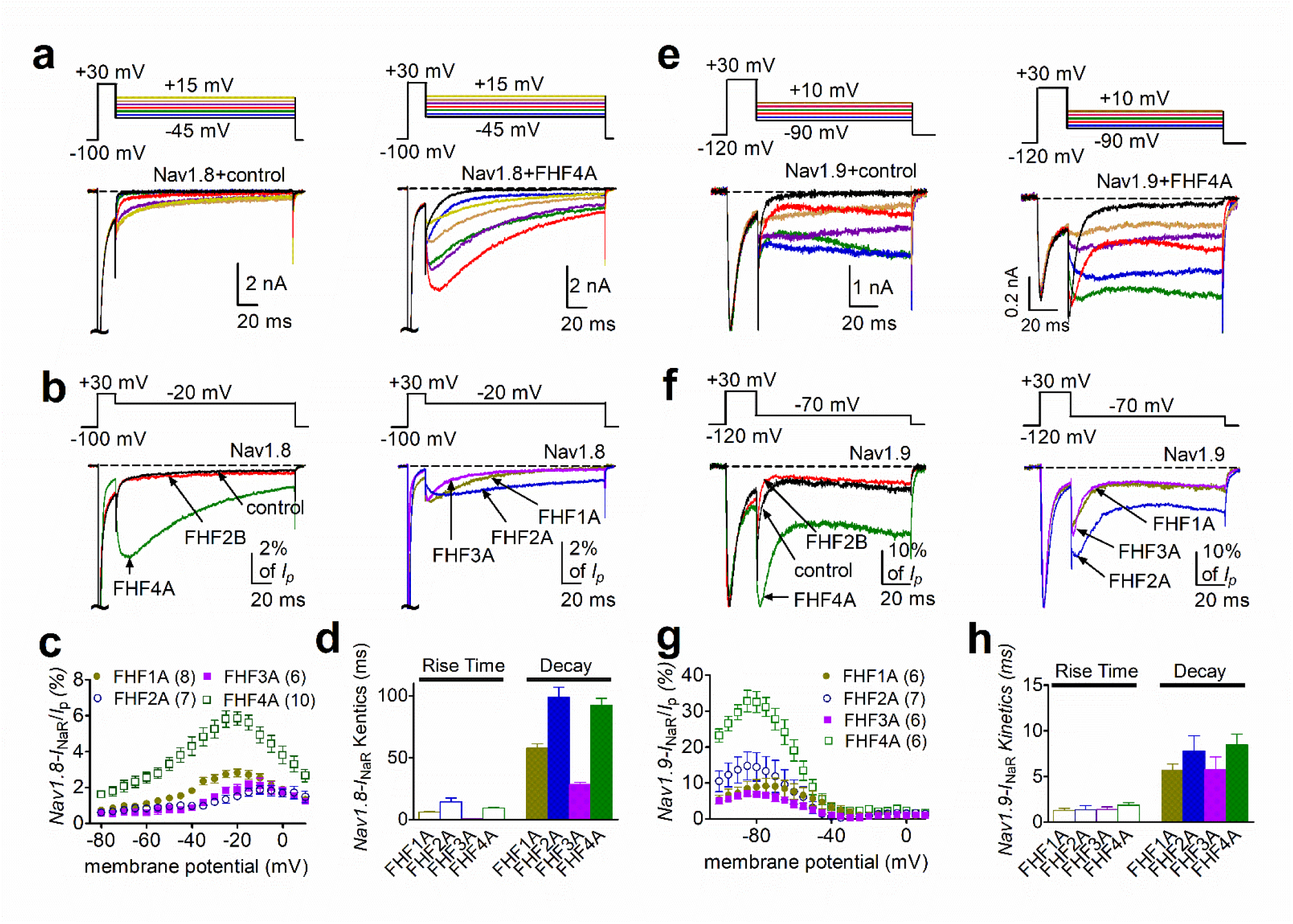
*I*_NaR_ were produced by recombinant Nav1.8 and Nav1.9 coexpressed with FHF2A or FHF4A in heterologous systems. (**a**, **e**) Family of representative current traces recorded from cells expressing Nav1.8 or Nav1.9that generated *I*_NaR_ in the presence of FHF4A (*right*) and that did not in the absence of any FHFs (control, *left*). Currents were elicited by a standard resurgent current protocol shown in the *inset*. (**b**,**f**) Overlay of single current traces of Nav1.8 – Nav1.9 elicited by the protocol (*inset*) in the absence (control, *black*) or presence of FHF2B (*red*), FHF1A (*yellow*), FHF2A (*blue*), FHF3A (*purple*) and FHF4A (*green*). (**c**,**g**) Voltage dependence of the relative Nav1.8 and Nav1.9 *I*_NaR_ mediated by FHF1A - FHF4A. (**d**,**h**) The rise time (*time to peak*) and time constants of the decay kinetics of FHF-mediated *I*_NaR_ in Nav1.8 and Nav1.9. While cells expressing Nav1.8 were held at -100 mV, cells expressing Nav1.9 were at -120 mV. The number of separate cells tested is indicated in parentheses. Data points are shown as mean ± S.E.

This non-decaying current activated extremely slowly during 100 ms voltage pulses, occurred in the absence and presence of FHFs, and persisted even when the repolarization pulse was extended to 1000 ms (Figure 2-figure supplement 1*a*). Importantly, the current-voltage curve almost completely overlapped that of a predicted “window current” formed by superimposition of steady-state activation and inactivation curves (Figure 2-figure supplement 1*b*-*d*). Since “window current” typically results in persistent current (Attwell et al., 1979), we suggest that these non-decaying currents result from a slow recovery from inactivation of Nav1.9 persistent currents. Regardless, these data indicate for the first time that Nav1.9 channels can generate a novel *I*_NaR_ distinct from those generated by Nav1.8 and TTX-sensitive VGSCs.

In addition to FHF2A and FHF4A, FHF1A and FHF3A are also A-type FHFs and are predominantly expressed in the CNS (Liu et al., 2001; Goetz et al., 2009). Because ectopic Nav1.8 expression has been observed in CNS neurons in multiple sclerosis (Black et al., 2000), we examined whether FHF1A and FHF3A might induce *I*_NaR_ in Nav1.8 and Nav1.9 as well. In addition to FHF2A and FHF4A, FHF1A and FHF3A both induced *I*_NaR_ in Nav1.8 (FHF1A, 2.8% ± 0.2%; FHF3A, 1.9% ± 0.3%) and Nav1.9 (FHF1A, 9.0% ± 2.3%; FHF3A, 5.8% ± 1.1%). These *I*_NaR_ displayed a voltage dependence of activation similar to those observed with FHF2A and FHF4A (Fig. 1*b-f*). Based on the relative amplitudes of the generated *I*_NaR_, the rank order of the ability of the four A-type FHFs to mediate Nav1.8 *I*_NaR_ is FHF4A > FHF1A > FHF3A ≈ FHF2A, while the rank order for Nav1.9 *I*_NaR_ generation is FHF4A > FHF2A > FHF1A > FHF3A.

### F2A/F4A peptides fully reconstituted *I*_NaR_

FHF2A, but not FHF2B, induces robust *I*_NaR_ from Nav1.8 and Nav1.9 (Fig. 2*b*,*f*). Intriguingly, FHF2A and FHF2B differ only in their N-terminus due to the alternative splicing of exon 1 (Fig. 3*a*). Moreover, a peptide derived from amino acids 2-21 of the FHF2A N-terminus has been shown to induce long-term inactivation of Nav1.6 channels (Dover et al., 2010). We hypothesized that the same region of the N-terminal tail is the critical molecular component necessary for *I*_NaR_ induction by A-type FHFs. To test this hypothesis, we intracellularly applied a 20-residue peptide (F2A or F4A), derived from the N-terminal residues 2-21 of FHF2A or FHF4A. We asked whether these peptides could reconstitute A-type FHF-mediated *I*_NaR_ observed with coexpression of full-length FHF2A/FHF4A (Fig. 3*a*). In the presence of 1 mM F2A or F4A, both Nav1.8 and Nav1.9 generated *I*_NaR_. The relative amplitudes were 1.5% ± 0.1% and 2.5% ± 0.4% in Nav1.8 at -15 mV (Fig. 3*b*, c), and 18.0% ± 3.1% and 10.3% ± 1.4% in Nav1.9 at -85 mV, respectively (Fig. 3*f*, g). The *I*_NaR_ retained the kinetics and voltage dependence of activation as observed with full-length FHF2A and FHF4A (Fig. 2*c*, g). On the other hand, both F2A and F4A significantly decreased the inactivation time constant of the transient currents of Nav1.8 and Nav1.9 evoked by a 20-ms pre-pulse to +30 mV (Fig. 3*d*, g), suggesting that both F2A and F4A serve as open channel blockers of Nav1.8 and Nav1.9. This is consistent with the previous reports that F2A induces open-channel block in Nav1.5 and Nav1.6 (Venkatesan et al., 2014; Dover et al., 2010).

**Figure 3.**
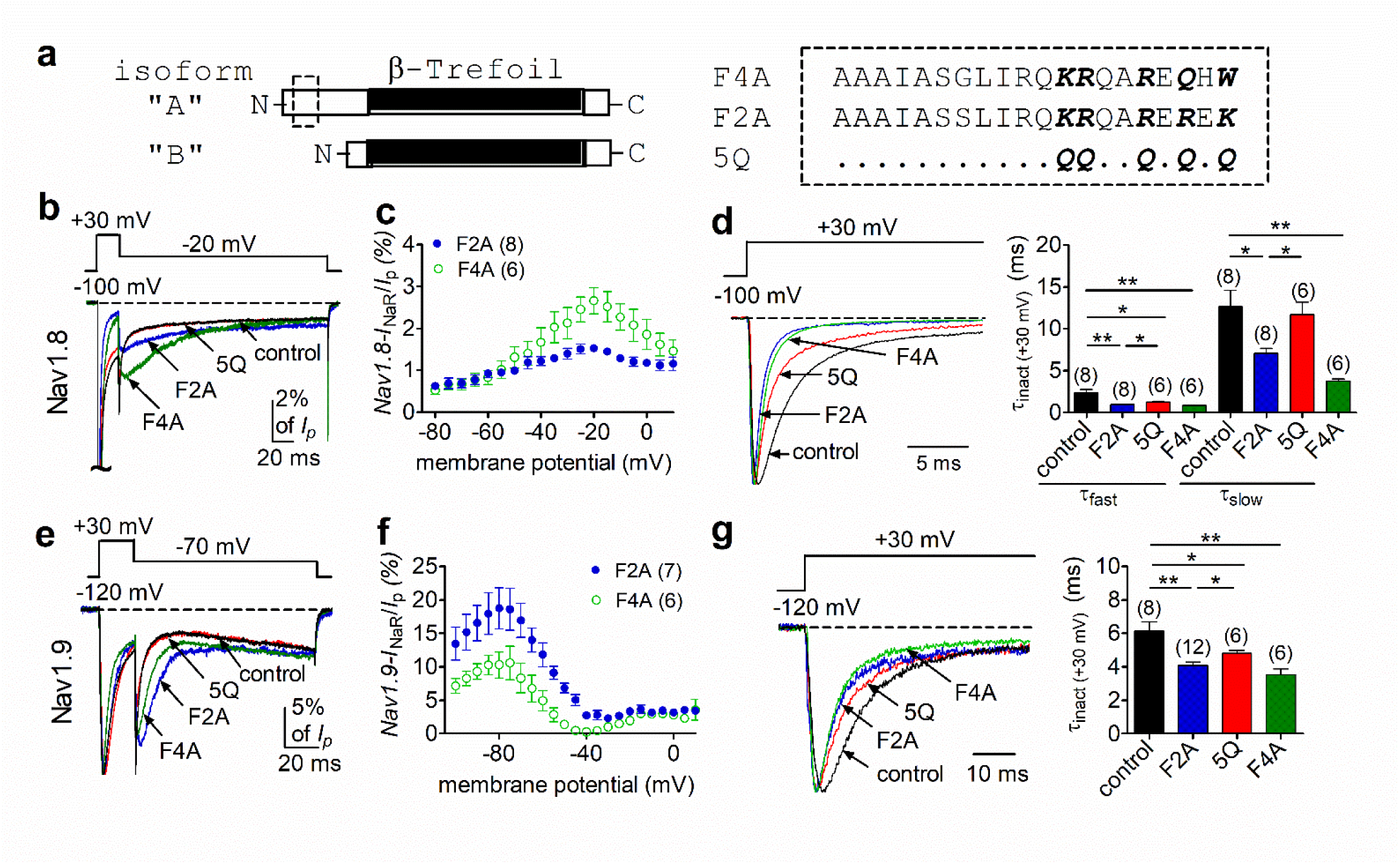
The peptides F2A and F4A fully reconstituted FHF2A/FHF4A-induced *I*_NaR_ in Nav1.8 and Nav1.9 in heterologous systems. (**a**) Schematic diagram of A- and B-type of FHFs (*left*). The amino acid sequences of short peptides located at N terminus of FHF2A and FHF4A are shown (*right*). Five positively charged residues of interest are highlighted in bold. 5Q is a mutant of F2A, in which five positive residues are replaced by Gln (Q). The residues conserved in F2A are indicated as dots. (**b**) Overlay of representative Nav1.8 *I*_NaR_ traces in the absence (control, black) and presence of F2A (blue), 5Q (red) or F4A (green). (**c**) Voltage dependence of the relative F2A- and F4A-induced Nav1.8 *I*_NaR_. (**d**) Decay time constants (τ, *right*) of transient Nav1.8 currents (*left*) at +30 mV. The time constants (τ_fast_, τ_slow_) were well fitted by a double exponential function. τ_fast_: control, 2.35 ± 0.40 ms; F2A, 0.92 ± 0.09 ms (p = 0.0037 *vs* control); 5Q, 1.26 ± 0.07 ms (p = 0.0339 *vs* F2A); F4A, 0.83 ± 0.04 ms (p = 0.0071 *vs* control). τ_slow_: control, 12.65 ± 1.95 ms; F2A, 7.07 ± 0.60 ms (p = 0.0161 *vs* control); 5Q, 11.67 ± 1.52 ms (p = 0.0065 *vs* F2A); F4A, 3.72 ± 0.28 ms (p = 0.0021 *vs* control). (**e**) Overlay of Nav1.9 *I*_NaR_ traces in the absence (control, black) and presence of F2A (blue), 5Q (red) or F4A (green). (**f**) Voltage dependence of the relative F2A- and F4A-induced Nav1.9 *I*_NaR_. (**g**) Decay time constants (τ, *right*) of transient Nav1.9 currents (*left*) at +30 mV. The time constants were fitted well by a single exponential function. Cells expressing Nav1.8 or Nav1.9 were held at -100 mV or -120 mV, respectively. All *I*_NaR_ of Nav1.8 or Nav1.9 were normalized to the peak transient current at -40 mV or at 0 mV, respectively. The concentrations of F2A, 5Q and F4A all are 1 mM. Filled and open circles represent FHF2A and FHF4A, respectively. The number of separate cells tested is indicated in parentheses. *, p < 0.05; **, p < 0.01.

To further investigate the roles of these peptides in *I*_NaR_ generation, we employed a F2A mutant (Dover et al., 2010) in which five positively charged residues (K1/R2/R3/R4/K5) are substituted with the neutral residue glutamine (5Q, Fig. 3*a*). In Fig. 3*b*,*e*, the mutant 5Q peptide failed to induce *I*_NaR_ in Nav1.8 or Nav1.9: the currents elicited during the repolarization pulse almost overlapped in the absence (control) and presence of 1 mM 5Q. Consistent with this finding, the transient current inactivated more slowly at +30 mV with 5Q than with F2A (Fig. 3*d*, g). Interestingly, the transient current still inactivated faster than under control conditions, suggesting that 5Q may still bind to VGSCs, but with lower affinity compared to F2A. These results suggest that the five positively charged residues in A-type FHFs are critical components for inducing *I*_NaR_.

### FHF4A mediated Nav1.8 *I*_NaR_ in sensory neurons

DRG neurons show expression of FHF4A along with Nav1.8 and Nav1.9. Our results in heterologous systems showed that FHF4A was most capable among the A-type FHF isoforms at inducing *I*_NaR_ with Nav1.8 and Nav1.9. Therefore, we next asked if FHF4A mediates *I*_NaR_ in primary neurons. We are able to isolate Nav1.9 *I*_NaR_ in DRG neurons (Fig. 4). However, while these unique *I*_NaR_ are strikingly similar to those recorded when A-type FHFs are co-expressed with Nav1.9 in HEK293 cells, the endogenous Nav1.9 *I*_NaR_ are only evident in a small subset of DRG neurons. On the other hand, in our previous work Nav1.8 type *I*_NaR_ could be recorded from the majority of DRG neurons expressing endogenous Nav1.8 currents and almost all DRG neurons expressing recombinant Nav1.8 (Xiao et al., 2019). We therefore next focused on interrogating the role of Nav1.8 *I*_NaR_ in DRG neurons.

**Figure 4.**
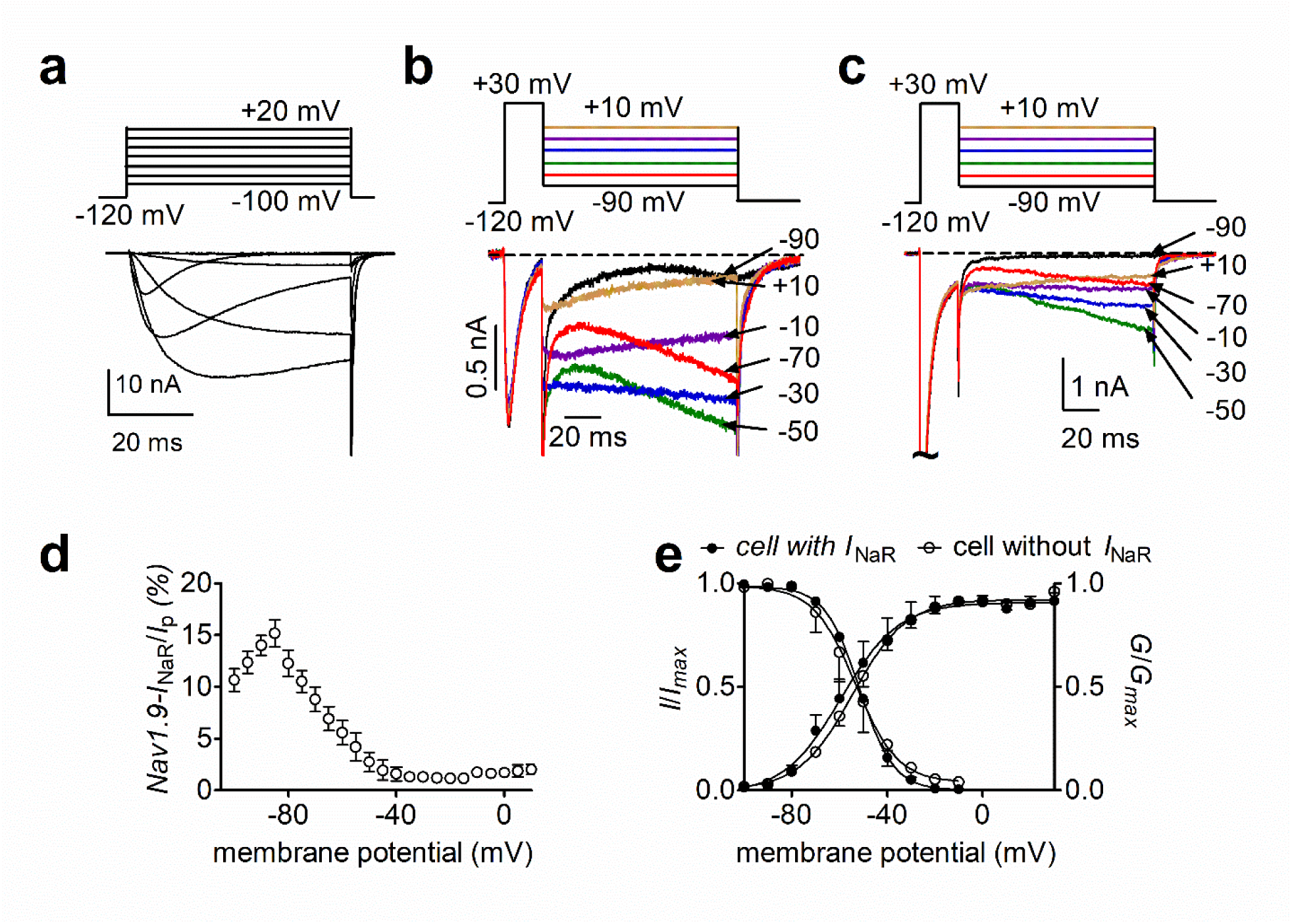
Nav1.9 *I*_NaR_ generated from Nav1.8 knockdown DRG neurons. (**a**), Typical Nav1.9 current traces induced by the protocol (*inset*), in which cells were subjected to 50-ms depolarization of potentials ranging from -120 to +40 mV with a 10-mV increment from a holding potential of -120 mV. (**b**,**c**), Representative current traces recorded from DRG neurons that did (**b**) and that did not (**c**) generate *I*_NaR_. Currents were elicited by a standard *I*_NaR_ protocol (*inset*), where cells were initially depolarized to +30 mV for 20 ms, then followed by a 100-ms hyperpolarizing potential ranging from +10 to –100 mV. (**d**), Voltage dependence of Nav1.9 *I*_NaR_ shown in (**b**). All *I*_NaR_ were normalized to peak transient current. (**e**), Steady-state activation and inactivation measured on DRG neurons with or without *I*_NaR_.

Consistent with our previous observation (Xiao et al., 2019), 13/14 small diameter DRG neurons transfected with a scrambled shRNA were found to generate Nav1.8 *I*_NaR_. The largest *I*_NaR_ was attained at -15 mV, with an average relative amplitude of 2.1% ± 0.3% of the peak transient TTX-resistant sodium current. The time to peak and the decay time constant for the current elicited at -15 mV were 45.0 ± 4.4 ms and 546.3 ± 43.2 ms, respectively. These results were identical to those seen in DRG neurons, without scrambled shRNA, in our previous work (Xiao et al., 2019), suggesting that the scrambled shRNA did not alter Nav1.8 *I*_NaR_. The efficiency of FHF4shRNA-mediated knockdown was determined using a monoclonal antibody specific to FHF4, which has been validated in heterologous systems in our laboratory. In Fig. 5*a,b*, FHF4shRNA reduced FHF4 expression by 73.1% (p < 0.0001). FHF4 knockdown did not significantly alter current density, voltage dependence of activation or recovery rate from inactivation of Nav1.8 currents in DRG neurons, but caused a hyperpolarizing shift of 12 mV in the voltage dependence of steady-state inactivation (p < 0.0001; Fig. 5*c-e*; Table 2). FHF4 knockdown considerably decreased the proportion of DRG neurons producing *I*_NaR_ (9/18 cells *vs* 13/14 scramble cells; p = 0.0095, χ^2^ test; Fig. 5*g*). Furthermore, in those DRG neurons with *I*_NaR_, FHF4 knockdown did not modify the voltage dependence of activation of Nav1.8 *I*_NaR_, but reduced the relative amplitude by about 42% (FHF4shRNA, 1.2% ± 0.2%; p < 0.05; Fig. 5*h*). Although our previous work showed that Navβ4 can contribute to generation of Nav1.8 *I*_NaR_ in DRG neurons (Xiao et al., 2019), the reduction here was Navβ4 independent because FHF4 knockdown did not significantly change Navβ4 expression level in our immunostaining experiments (Figure 5-figure supplement 1). Therefore, our data indicate that FHF4A is a major producer of Nav1.8 *I*_NaR_ in small diameter DRG neurons. The remaining *I*_NaR_ after FHF4-knockdown are possibly mediated by residual FHF4A, endogenous FHF2A or endogenous Navβ4.

**Figure 5.**
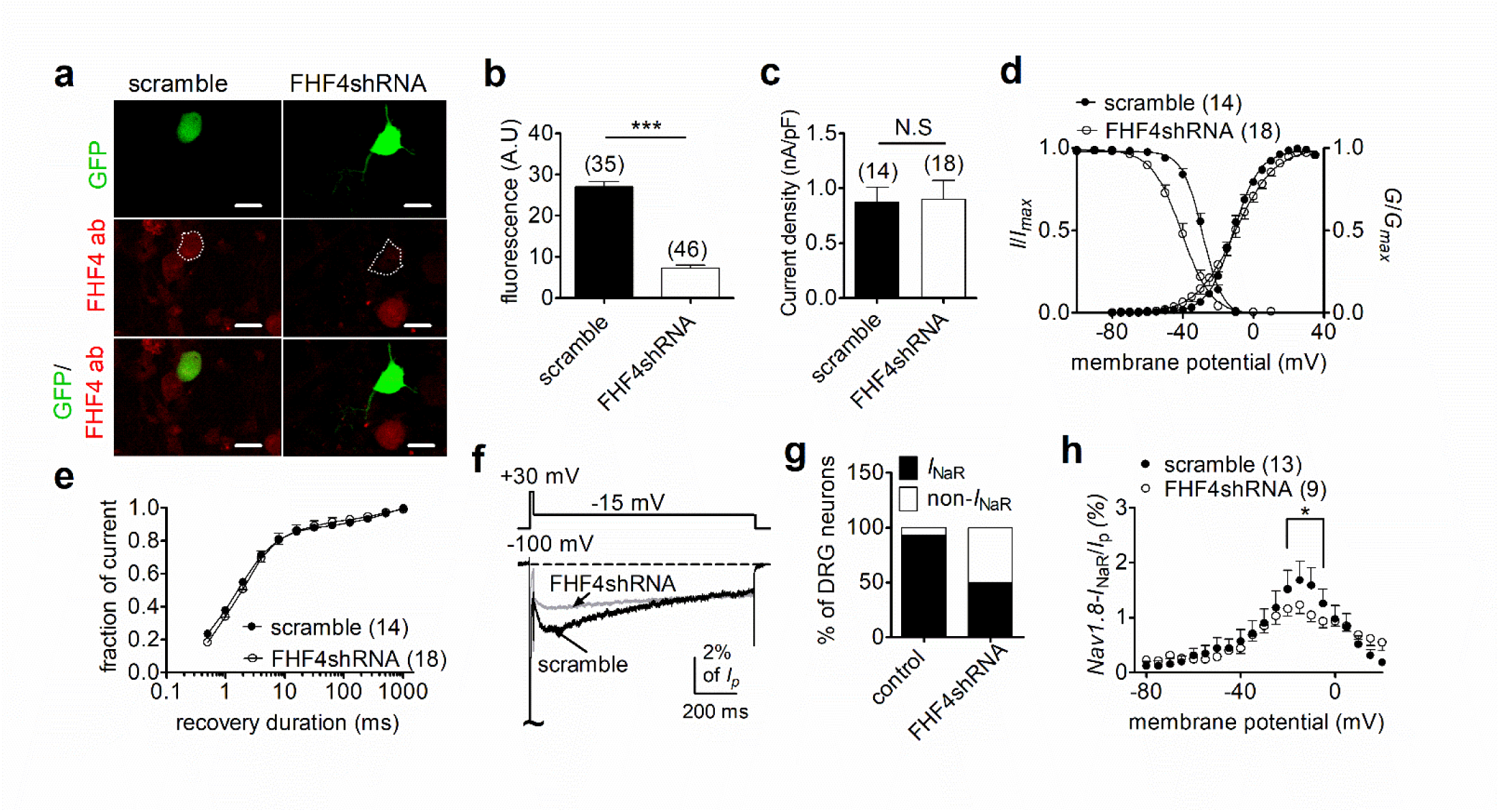
FHF4 knockdown reduced the ability of Nav1.8 to generate *I*_NaR_ in rat DRG neurons. (**a**) Immunofluorescent reactions showed expression levels of FHF4 in DRG neurons. Dashed lines show the shape of transfected DRG neurons. Scale bars, 50 µm; ab, antibody. (**b**) Summary of fluorescence in DRG neurons transfected with the scrambled shRNA or FHF4shRNA (p < 0.0001). (**c**) FHF4 knockdown did not significantly alter Nav1.8 current density (p = 0.9116). (**d**) FHF4 knockdown shifted voltage dependence of steady-state inactivation to more negative potentials (p < 0.0001), but did not affect activation (p = 0.9116). (**e**) FHF4 knockdown did not significantly impair the recovery rate from inactivation The time constants estimated from single exponential fits were 2.92 ± 0.53 ms (scramble) and 4.00 ± 1.01 ms (FHF4shRNA, p = 0.3905), . (**f**) *I*_NaR_ traces recorded from small diameter DRG neurons transfected with scramble or FHF4shRNA. (**g**) FHF4 knockdown decreased the percentage of DRG neurons to generate Nav1.8 *I*_NaR_ (p < 0.0001). (**h**) Voltage dependence of the relative Nav1.8 *I*_NaR_ in DRG neurons treated with scramble and FHF4shRNA. Filled and open circles represent scramble and FHF4shRNA, respectively. The number of separate cells tested is indicated in parentheses. N.S, not significant; *, p < 0.05; ***, p < 0.0001.

**Table 2.** Gating properties of Nav1.8 in DRG neurons

| $V_{1/2}$ | Control (mV) | FHF4shRNA | +F4A |
| --- | --- | --- | --- |
| activation | $-11.1 \pm 1.1$ (14) | $-10.7 \pm 2.9$ (18) | $-9.3 \pm 3.5$ (10) |
| inactivation | $-28.5 \pm 1.0$ (14) | $-40.4 \pm 2.0^{\text{a}}$ (18) | $-35.9 \pm 1.2^{\text{a}}$ (10) |

### FHF4A-mediated Nav1.8 *I*_NaR_ regulated sensory neuron excitability

We next explored the impact of FHF4A-mediated *I*_NaR_ on neuronal excitability. Previous studies have shown that FHFs profoundly modulate the activities of TTX-sensitive VGSCs in DRG neuron (Barbosa et al.; 2017, Venkatesan et al., 2014; Dover et al., 2010), therefore we measured excitability of small diameter DRG neurons in the presence of 500 nM TTX, which blocks all TTX-sensitive VGSCs and removes this confounding variable. FHF4 knockdown did not change resting membrane potential, input resistance or current threshold of action potential firing (scramble, -54.9 ± 1.1 mV, 463.3 ± 38.5 MΩ, 1.26 ± 0.21 nA; FHF4shRNA, -57.2 ± 1.6 mV, 548.7 ± 66.9 MΩ, 1.22 ± 0.14 nA; p > 0.05; Fig. 6*a*-*c*), but narrowed single evoked action potentials. The average action potential durations measured under scramble and FHF4shRNA were 17.94 ± 2.63 ms and 10.54 ± 1.19 ms (p = 0.0153; Fig. 6*d*), respectively. With 2-s injected currents greater than 300 pA, the FHF4-knockdown DRG neurons displayed significantly fewer action potentials than the scramble-treated neurons (Fig. 6*e*, *f*).

**Figure 6.**
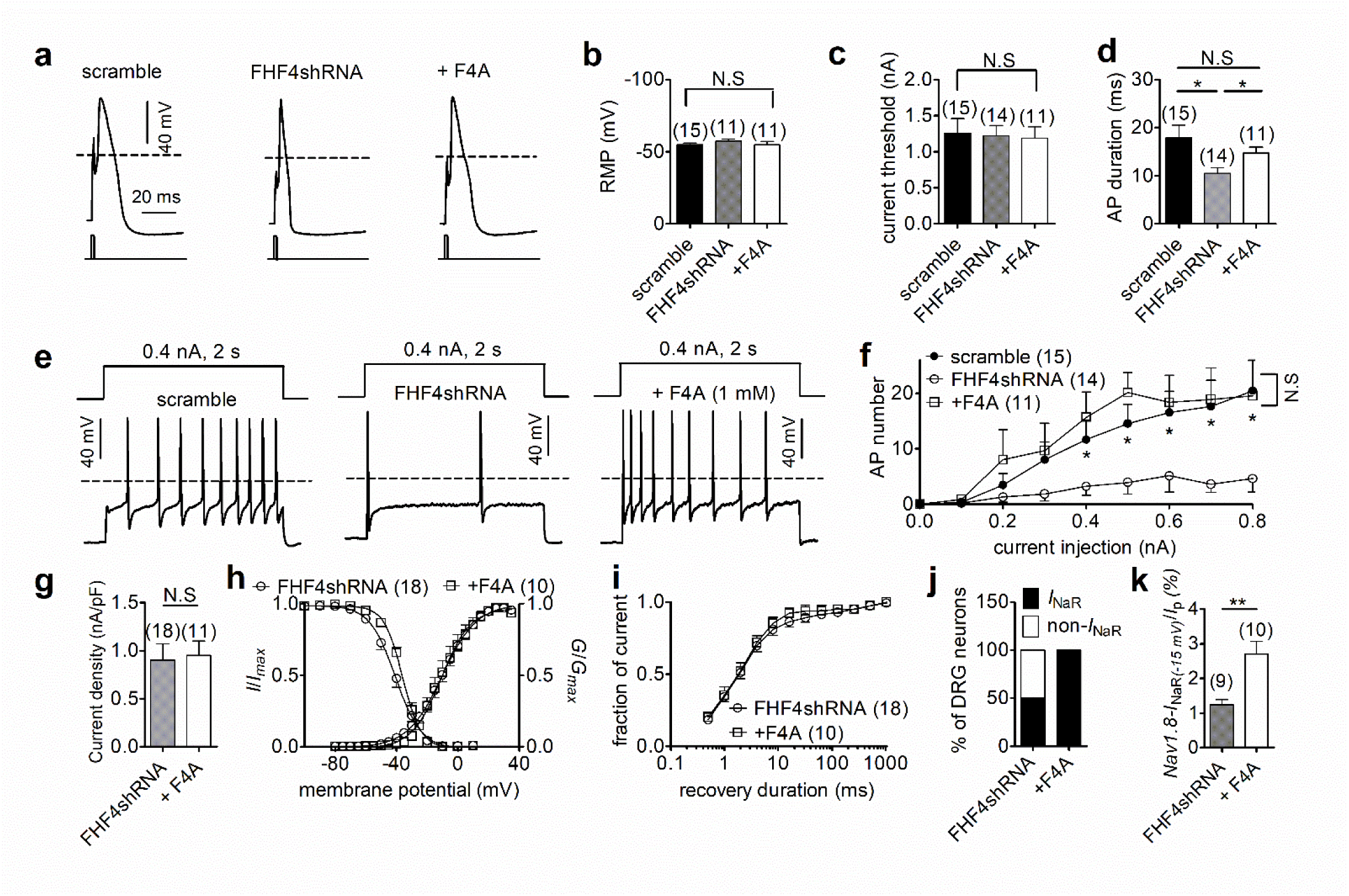
FHF4shRNA-mediated reduction in DRG neuron excitability was rescued by the F4A peptide. (**a**) Typical single action potentials elicited by a 1-ms current injection. (**b**) Resting membrane potentials under scramble, FHF4shRNA and FHF4shRNA + F4A (p = 0.6149, one-way ANOVA). (**c**) Summary of current threshold (p = 0.9673, one-way ANOVA). (**d**) Summary of action potential duration (APD90). The durations were 17.94 ± 2.63 ms (scramble), 10.54 ± 1.19 ms (FHF4shRNA, p = 0.0153 *vs* control) and 14.76 ± 1.22 ms (+F4A, p = 0.0233 *vs* FHF4shRNA and p = 0.3011 *vs* control), respectively. (**e**) Typical action potential trains elicited by a 2-s injection of 400-pA current. (**f**) Summary of the number of action potentials elicited by a 2-s injection of currents ranging from 0 – 800 pA. (**g**) F4A did not alter Nav1.8 current density (p = 0.8428). (**h**) Voltage dependence of activation and steady-state inactivation of Nav1.8 before and after addition of F4A in FHF4shRNA-treated DRG neurons (activation: p = 0.8160; inactivation: p = 0.0332). (**i**) F4A did not impair the recovery rate from Nav1.8 inactivation in FHF4shRNA-treated DRG neurons. The time constants estimated from single exponential fits were 4.00 ± 1.01 ms (FHF4shRNA) and 2.92 ± 0.42 ms (+F4A, p = 0.5826), respectively. (**j**) F4A increased the percentage of FHF4shRNA-treated DRG neurons to generate Nav1.8 *I*_NaR_ (p = 0.0066). (**k**) F4A increased the relative amplitude of Nav1.8 *I*_NaR_ in FHF4shRNA-treated DRG neurons (p = 0.0027). In (**a**–**k**), the concentration of F4A is 1 mM. Filled circles, open circles and open squares represent scramble, FHF4shRNA and F4A, respectively. The number of separate cells tested is indicated in parentheses. The V_1/2_ values measured in (**h**) were summarized in Table 2. N.S, not significant; *, p < 0.05; ***, p < 0.001.

We then tested whether F4A peptide could reverse the FHF4A knockdown-mediated effects on *I*_NaR_ and neuronal excitability. Intracellular application of 1 mM F4A did not significantly alter current density, voltage dependence of activation, steady-state inactivation or recovery rate from inactivation of Nav1.8 currents in FHF4shRNA-treated DRG neurons (Fig. 6*g*-*i*; Table 2). Although F4A peptide did not reverse the negative shift in the voltage dependence of steady-state inactivation caused by FHF4 knockdown (shown in Fig. 5*d*), F4A peptide did rescue the FHF4-knockdown-mediated decrease in *I*_NaR_: 10/10 DRG neurons tested yielded Nav1.8 *I*_NaR_ (p = 0.0066; χ^2^ test; Fig. 6*j*). The average relative amplitude measured at -15 mV increased from 1.2% ± 0.2% (FHF4shRNA) to 2.7% ± 0.4% (F4A; p = 0.0027; Fig. 6*k*), similar to the amplitude yielded under the scramble condition. F4A peptide did not change the resting membrane potential, input resistance or current threshold in FHF4shRNA-treated DRG neurons (+F4A, - 54.8 ± 2.3 mV, 502.8 ± 92.9 MΩ, 1.19 ± 0.15 nA; p > 0.05 *vs* scramble and FHF4shRNA; Fig. 6*b*, *c*), but significantly broadened action potentials (average duration of 14.76 ± 1.22 ms; p = 0.0233; Fig. 6*d*). F4A peptide increased the number of action potentials elicited by 2-s injected currents of 400 pA (Fig. 6*e*). Finally, the FHF4shRNA-transfected DRG neurons treated with F4A peptide could fire action potentials at almost the same frequency as the scramble-transfected neurons (Fig. 6*f*), demonstrating that the loss of sensory neuron excitability by FHF4 knockdown can be rescued by F4A. Therefore, our data clearly illustrate that A-type FHF is a critical molecule in small diameter DRG neurons and that A-type FHF determines neuronal excitability via *I*_NaR_ generation.

### Navβ4 does not elicit Nav1.9 *I*_NaR_

The Nav1.9 *I*_NaR_ identified in HEK293 cells cotransfected with Nav1.9 and A-type FHFs is distinct from the *I*_NaR_ observed with other VGSCs. As Navβ4 has been shown to induce *I*_NaR_ in all of the other VGSC isoforms (Nav1.1 – Nav1.8), we asked if Navβ4 also induces *I*_NaR_ with Nav1.9. Multiple studies have failed to reconstitute *I*_NaR_ in heterologous systems by coexpressing full-length Navβ4 with VGSC α-subunits (12, 39, 40). However, a short peptide (KKLITFILKKTREK) derived from the Navβ4 C-terminus can induce *I*_NaR_ generation in heterologous expression systems and also in primary neurons (Grieco et al., 2005; Barbosa et al., 2015). Here we intracellularly applied Navβ4 peptide (200 µM) to investigate if Nav1.9 expressed in HEK293 cells could utilize the C-terminus of Navβ4 to generate *I*_NaR_. Surprisingly, Navβ4 peptide did not induce Nav1.9 *I*_NaR_. Only classic tail currents were observed with the Navβ4 peptide (p > 0.05; Fig. 7*a*, *b*), indicating that Navβ4 is not capable of mediating *I*_NaR_ in Nav1.9.

**Figure 7.**
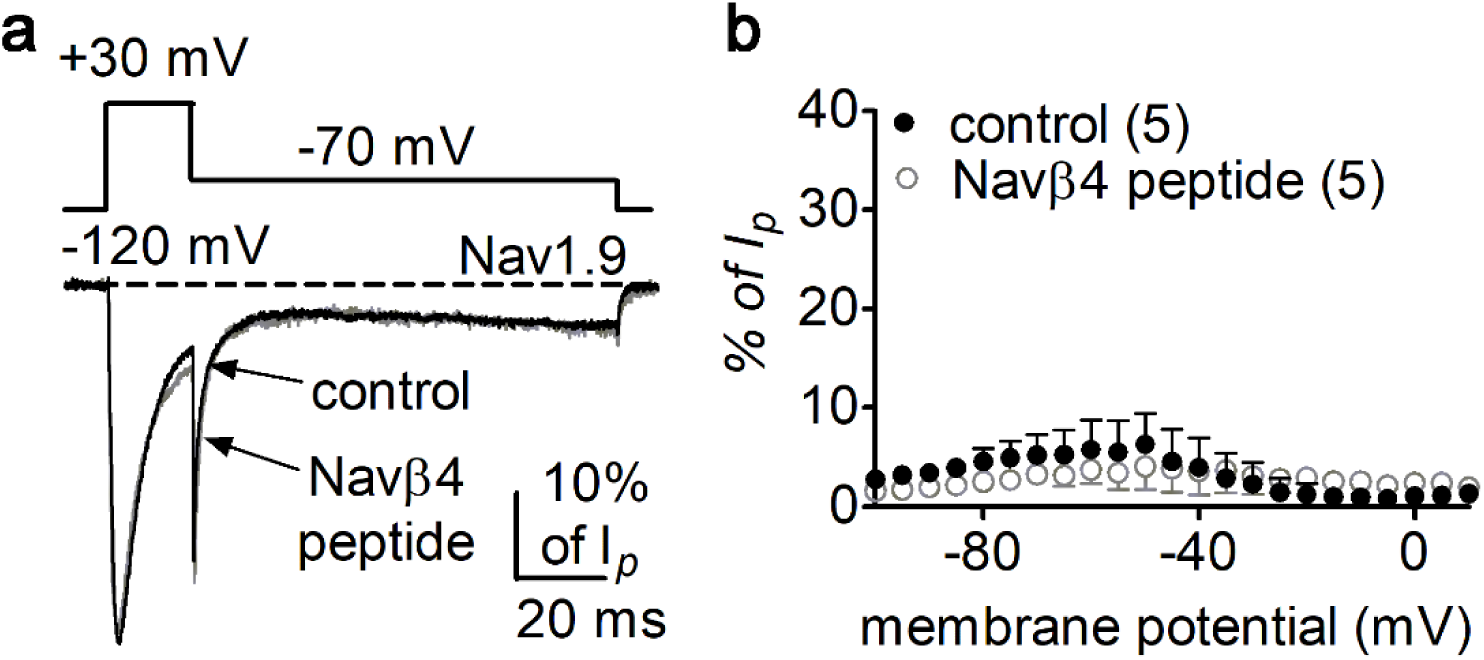
Navβ4 peptide did not induce Nav1.9 *I*_NaR_ in HEK293 cells. (**a**) Overlay of normalized current traces elicited by a resurgent protocol (*inset*) in the absence (control, *grey*) and presence of 200 µM Navβ4 peptide (*black*). (**b**) Voltage dependence of the relative currents. Filled and open circles represent control and Navβ4 peptide, respectively.

### An inner pore residue impairs Navβ4, but not A-type FHFs, *I*_NaR_

Nav1.9 exhibits low (42-53%) sequence similarity to other mammalian VGSC subtypes. We hypothesized that non-conserved pore residues, especially positive residues, might prevent the positively charged Navβ4 peptide from binding to the Nav1.9 inner pore. Sequence analysis identified K799, residing in the II-S6 segment of Nav1.9, as a promising candidate for such prevention (Fig. 8*a*). The corresponding residue in all other VGSC isoforms is an asparagine. As described previously (Lin et al., 2016), the K799N mutation does not significantly alter Nav1.9 gating properties (Fig. 8*b*; Table 3). Interestingly, the K799N mutation greatly enhanced the ability of Navβ4 peptide (200 µM) to mediate Nav1.9 *I*_NaR_ in response to a depolarizing voltage of +100 mV. Fig. 8*c* shows that Navβ4 peptide mediated *I*_NaR_ in K799N channels with a fast onset/decay kinetics and a hyperpolarized voltage dependence of activation similar to A-type FHF-mediated Nav1.9 *I*_NaR_. The relative amplitude is 6.6% ± 0.5% at -85 mV (Fig. 8*d*). Intriguingly, the K799N mutation did not alter F2A-mediated *I*_NaR_ (Nav1.9, 17.0% ± 2.5%; K799N, 18.1% ± 3.5%; p > 0.05; Fig. 8*e, f*).

**Figure 8.**
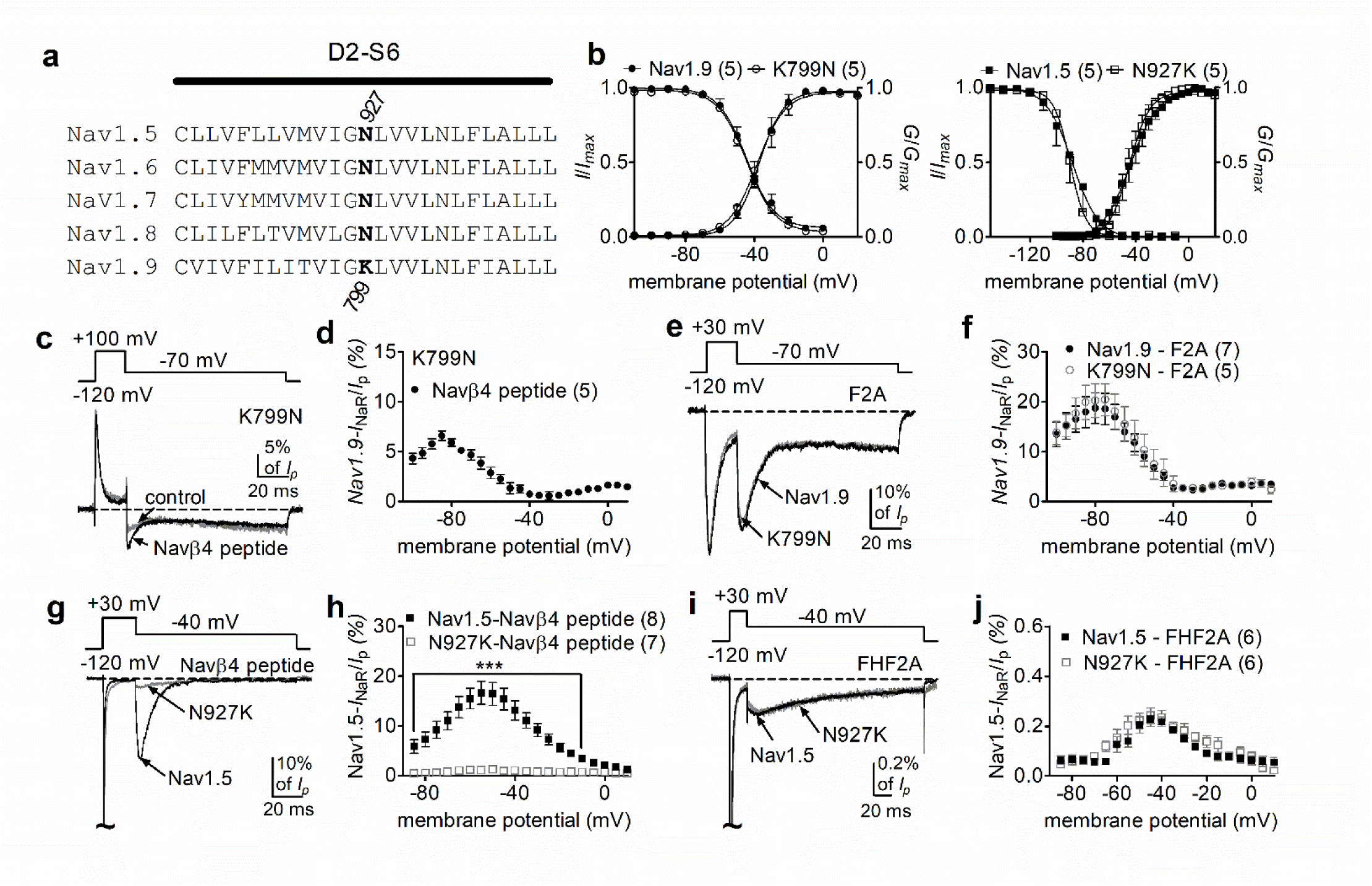
The residue at position 799 in Nav1.9 was crucial for VGSC sensitivity to Navβ4. (**a**) Sequence alignment of domain II S6 segments of Nav1.5 - Nav1.9. The position of the residues of interest is indicated in bold and designated with a number. (**b**) The K799N mutation and the reversal mutation N927K did not significantly alter steady-state activation or inactivation of Nav1.9 (circles, *right*) and Nav1.5 (squares, *left*), respectively. (**c**) The Nav1.9 mutant K799N generated *I*_NaR_ in the presence of 200 µM Navβ4 peptide (*black*). Control, *grey*. (**d**) Voltage dependence of the relative *I*_NaR_ in the Nav1.9 mutant K799N (filled circles). (**e**) Typical *I*_NaR_ traces recorded from Nav1.9 (*black*) and the mutant K799N (*grey*) in the presence of 1 mM F2A. (**f**) Comparison of the relative F2A-induced *I*_NaR_. Filled and open circles represent Nav1.9 and the mutant K799N, respectively. (**g**) Typical *I*_NaR_ traces recorded from Nav1.5 (*black*) and the mutant N927K (*grey*) in the presence of 200 µM Navβ4 peptide. (**h**) Voltage dependence of the relative *I*_NaR_ in Nav1.5 (filled squares) and the mutant N927K (open squares). (**i**) Typical *I*_NaR_ traces recorded from Nav1.5 (*black*) and the mutant N927K (*grey*) in the presence of FHF2A. (**j**) Comparison of the relative FHF2A-induced *I*_NaR_ in Nav1.5 (filled squares) and the mutant N927K (open squares). In (**c,e**,**g**,**i**), *I*_NaR_ were elicited by the protocols shown in the *inset*. In (**b**,**c**,**d**,**g**,**h**), 500 µM GTP-γ-S was added for Nav1.9 and K799N cells in the pipette solution. F2A and Navβ4 peptide were applied in peptide solution. The number of separate cells tested is indicated in parentheses. ***, p < 0.005.

**Table 3.** Gating properties of wild-type Nav1.5, the mutant N927K, wild-type Nav1.9 and the mutant K799N.

| $V_{1/2}$ (mV) | Nav1.5wt | N927K | Nav1.9wt | K799N |
| --- | --- | --- | --- | --- |
| activation | $-44.6 \pm 4.4$ (5) | $-44.0 \pm 2.9$ (5) | $-37.1 \pm 3.3$ (5) | $-37.5 \pm 4.1$ (5) |
| inactivation | $-88.5 \pm 6.4$ (5) | $-89.7 \pm 0.9$ (5) | $-45.7 \pm 1.9$ (7) | $-44.0 \pm 1.5$ (5) |

To further confirm the role of this residue in modulating VGSCs *I*_NaR_, we constructed reverse mutations at corresponding positions in Nav1.5 and Nav1.7 (Fig. 8*a*). The reverse mutation N927K did not influence gating properties of Nav1.5 (Fig. 8*b*; Table 3), but reduced Nav1.5 *I*_NaR_ induced by the presence of 200 µM Navβ4 peptide by 92% (Nav1.5, 17.2% ± 2.1%; N927K, 1.3% ± 0.1%; P < 0.0001; Fig. 8*g*, *h*). A substantial reduction (∼85%) was also observed for the N945K mutation in Nav1.7 (Figure 8-figure supplement 1). In contrast, the N927K mutation did not impair the ability of Nav1.5 to generate *I*_NaR_ mediated by full-length FHF2A (Nav1.5, 0.24% ± 0.03%; N927K, 0.23% ± 0.03%; p > 0.05; Fig. 8*i*, *j*). Collectively, these results indicate that the residue at position 799 in Nav1.9 is involved in VGSC interaction with Navβ4. Although K799 in Nav1.9 is a major determinant of Nav1.9 resistance to the Navβ4 peptide, it may not be the only factor involved in Nav1.9 resistance. Furthermore, because changes at this position did alter A-type FHF mediated *I*_NaR_, we propose that A-type FHFs and Navβ4 do not share identical binding determinants in the pore of VGSCs.

### FHF2A mediated Nav1.5 and Nav1.7, but not Nav1.6, *I*_NaR_ in heterologous system

Finally, we asked if other VGSC isoforms share the FHF mechanism of *I*_NaR_ generation. We studied Nav1.5, Nav1.6 and Nav1.7, because they are coexpressed with FHF2A or FHF4A in cardiac myocytes and neurons (Li et al., 2002; Wang et al., 2011a; Yan et al., 2014; Barbosa et al., 2017; White et al., 2019). Coexpression of FHF2A with Nav1.5 or Nav1.7 induced *I*_NaR_ (Fig. 9*a-f)*, with a voltage dependence of activation more negative than Nav1.8 but more positive than Nav1.9 *I*_NaR_ (Fig. 1*c*, *f*). Maximal *I*_NaR_ were attained at near - 40 mV. The average relative amplitudes were at least 6-fold smaller (Nav1.5, 0.22% ± 0.02%; Nav1.7, 0.27% ± 0.02%) than Nav1.8 and Nav1.9 *I*_NaR_. The time to peak and the decay time constant for the FHF2A-mediated Nav1.5 *I*_NaR_ were 10.2 ± 1.3 ms and 120.8 ± 19.6 ms at -40 mV, respectively (Fig. 9*i*). FHF2A-mediated Nav1.7 *I*_NaR_ displayed a similar rise and decay kinetics to Nav1.5 *I*_NaR_ (Fig. 9*i*).

**Figure 9.**
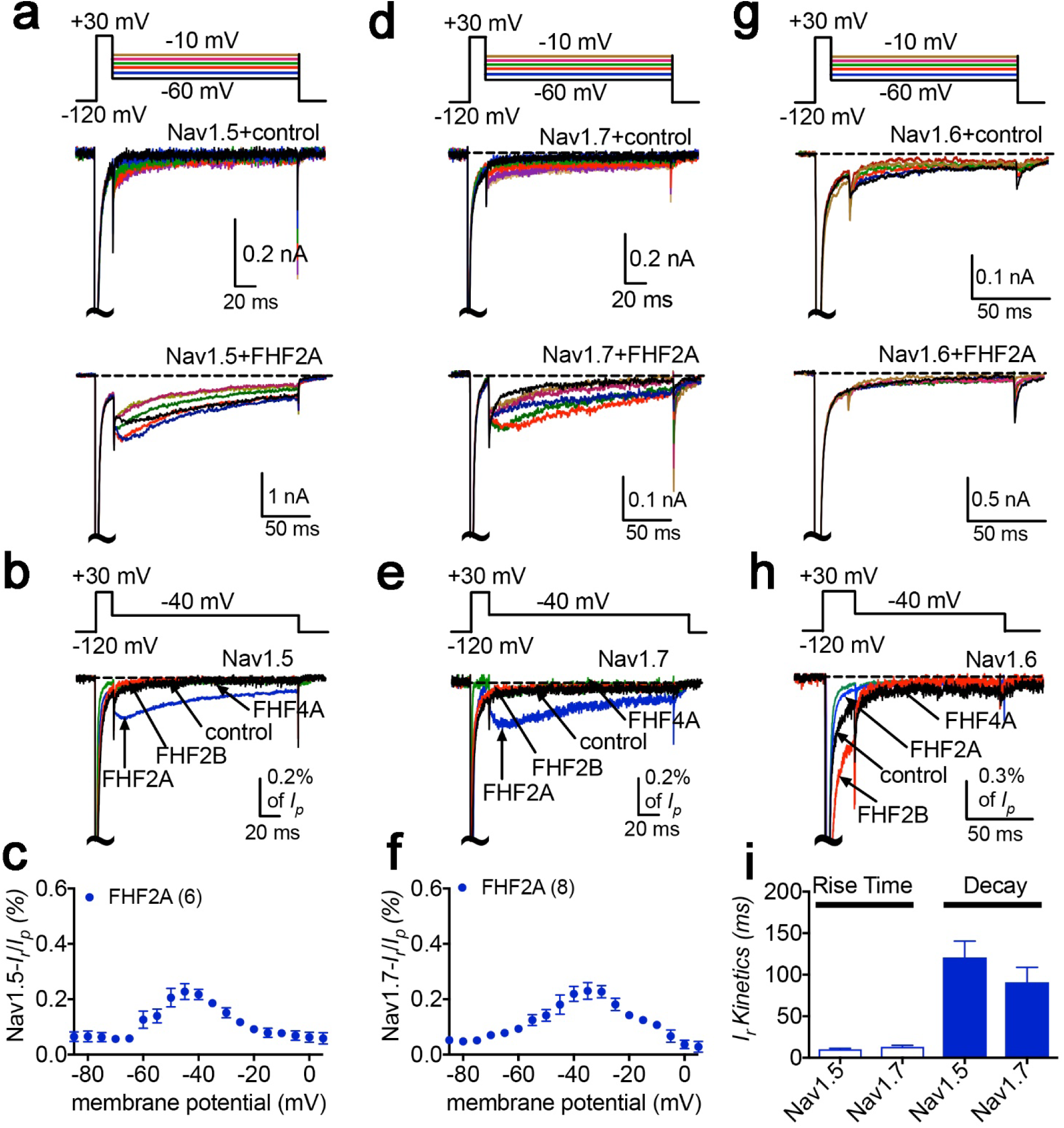
*I*_NaR_ were produced by recombinant Nav1.5 and Nav1.7 coexpressed with FHF2A in heterologous systems. (**a**,**d**,**e**) Family of representative current traces recorded from cells expressing Nav1.5, Nav1.7, or Nav1.6 in the presence of FHF2A (*below*) and that did not in the absence of any FHFs (control, *upper*). Currents were elicited by a standard *I*_NaR_ protocol shown in the *inset*. (**b**,**e**,**h**) Overlay of single current traces of Nav1.5 – Nav1.7 elicited by the protocol (*inset*) in the absence (control, *black*) or presence of FHF2B (*red*), FHF2A (*blue*) and FHF4A (*green*). (**c**,**f**) Voltage dependence of the relative Nav1.5 and Nav1.7 *I*_NaR_ mediated by FHF2A. (**i**) The rise time and time constants of the decay kinetics of FHF2A-mediated *I*_NaR_ in Nav1.5 and Nav1.7. In (**c**,**f**), all *I*_NaR_ were normalized to the peak transient current. In (**i**), time constants were obtained by fitting a single exponential function. Cells were held at - 120 mV. The number of separate cells tested is indicated in parentheses. Data points are shown as mean ± S.E.

However, neither FHF2B nor FHF4A generated *I*_NaR_ in Nav1.5 and Nav1.7 (Fig. 9*b*, *e*). Neither FHF2A nor FHF4A induced generation of *I*_NaR_ with Nav1.6 (Fig. 8*g*, *h).* However, coexpression of FHF4A with Nav1.6 elicited long-term inactivation of Nav1.6 in ND7/23 cells (Figure 10), similar to that previously shown for co-expression of FHF2A with Nav1.6 in HEK293 cells (Rush et al., 2006). Furthermore, intracellular application of the F4A peptide did not induce *I*_NaR_, only long-term inactivation similar to that induced by full-length FHF4A (Figure 10).

**Figure 10.**
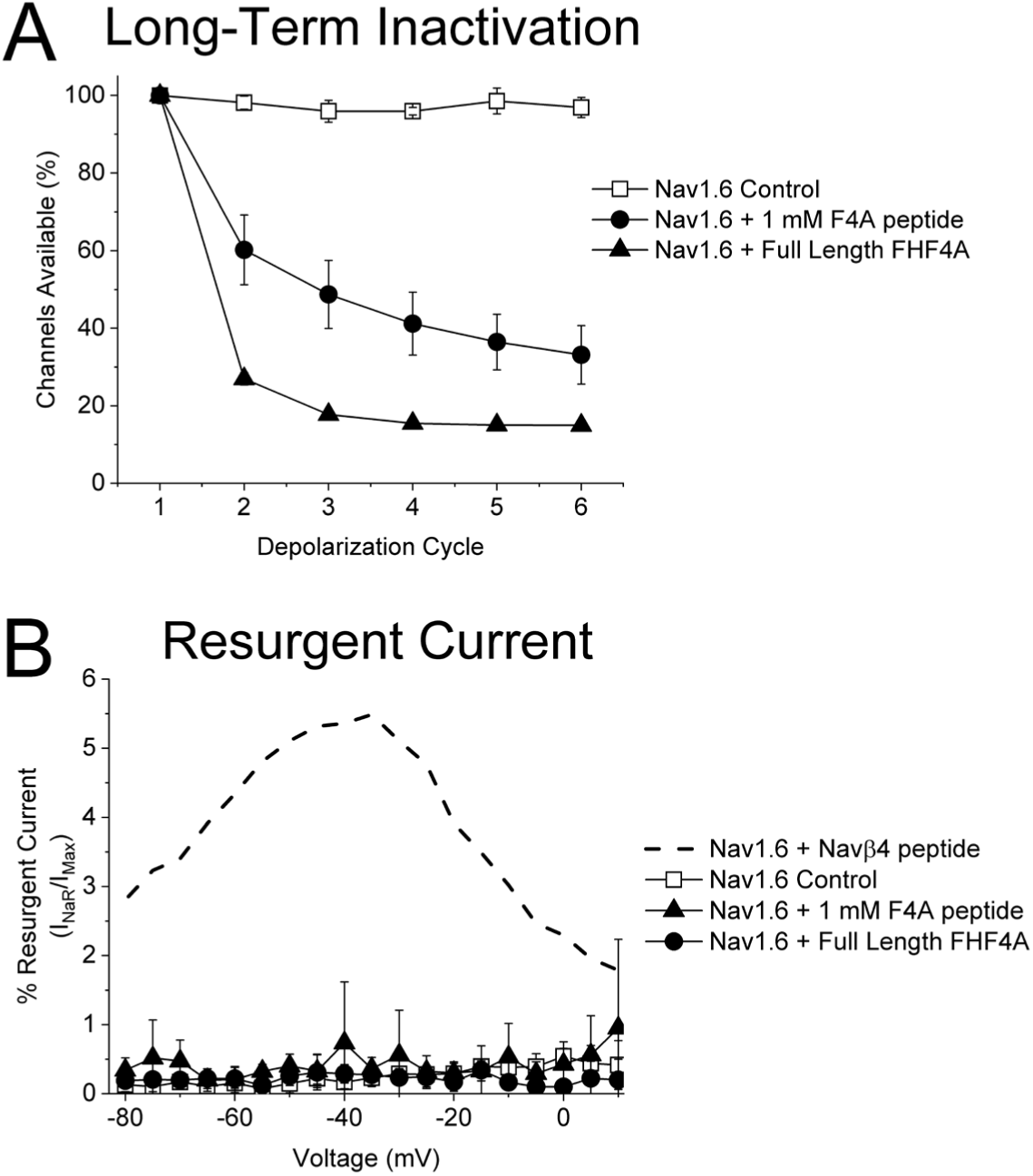
FHF4A induces long-term inactivation, not *I*_NaR_, in Nav1.6 channels. HEK293 cells stably expressing human Nav1.6 were recorded under control conditions, after FHF4A transfection and with F4A peptide (1 mM) in the pipette solution. (**a**) Both the full-length FHF4A and the F4A peptide induced a substantial increase in long-term inactivation in response to a train of six -20 mV depolarizations at ∼50 Hz. (**b**) Neither full-length FHF4A nor F4A peptide induced detectable *I*_NaR_ in HEK293 cells expressing Nav1.6 channels. For comparison, data for Nav1.6 *I*_NaR_ with Navβ4 peptide (200 mM) is shown with the dashed curve, adapted from Pan and Cummins (2020).

## Discussion

Resurgent sodium currents are critical regulators of central and peripheral neuron excitability. In this study, we identify A-type FHFs as direct mediators of TTX-resistant VGSC *I*_NaR_. We show, for the first time, that coexpression of only two proteins, a full-length A-type FHF and a VGSC α-subunit, is sufficient in heterologous systems to reconstitute *I*_NaR_. We show that short peptides derived from A-type FHF N-termini, the precise residues that induce long-term inactivation in other VGSCs (Venkatesan et al., 2014) can fully replicate the Nav1.8 and Nav1.9 TTX-resistant *I*_NaR_. Importantly, we implicate A-type FHFs as major drivers of TTX-resistant *I*_NaR_ in nociceptive DRG neurons.

We also identified a novel TTX-resistant Nav1.9 *I*_NaR_, which shows unique biophysical properties. The voltage dependence of activation of Nav1.9 *I*_NaR_ is >40 mV more negative than previously described *I*_NaR_ and these Nav1.9 currents exhibit faster onset/decay kinetics than Nav1.8 *I*_NaR_. The ratio of *I*_NaR_ to peak transient current is at least fivefold larger in Nav1.9 than in other VGSC isoforms. This is likely due to the extremely slow fast-inactivation of Nav1.9, because destabilizing VGSC fast-inactivation by disease mutations or toxins enhances *I*_NaR_ generation (Grieco and Raman, 2004; Jarecki et al., 2010; Bant et al., 2013; Xiao et al., 2019). Interestingly, while coexpression of A-types FHFs resulted in consistent Nav1.9 *I*_NaR_ in HEK293 cells, only a small fraction of rodent DRG neurons exhibited Nav1.9 *I*_NaR_. This might due to differences between rodent and human Nav1.9, or as a result of the complex modulation of Nav1.9 *I*_NaR_ in sensory neurons.

Our data reveal a novel mechanism of *I*_NaR_ generation independent of Navβ4. This is surprising, because a substantial number of studies have implicated Navβ4 as a major determinant of *I*_NaR_ in multiple VGSC subtypes (Grieco et al., 2005; Jarecki et al., 2010; Bant and Raman, 2010; Lewis and Raman, 2011; Miyazaki et al., 2014; Barbosa et al., 2015; Patel et al., 2016). Our mutagenesis experiments show that an inner pore residue (K799) that is unique to Nav1.9 determines the inability of Navβ4 peptide to induce Nav1.9 *I*_NaR_. The K799 residue is replaced by asparagine at the corresponding position (D2-S6) in Nav1.1 – Nav1.8. The homology model of Nav1.9 shows that the side chain of K799 projects into the channel pore (Lin et al., 2016), and thus the positively charged residue may prevent the positively charged Navβ4 peptide from accessing its binding site in the Nav1.9 inner pore by electrostatic repulsion, which is consistent with previous findings that positive residues are crucial for Navβ4 peptide mediating *I*_NaR_ (Lewis and Raman, 2011).

The most salient finding from this study is identification of A-type FHFs as novel *I*_NaR_ mediators of Nav1.8 and Nav1.9 and, to a lesser extent, Nav1.5 and Nav1.7. A-type FHFs consist of a long N-terminus, an FGF-like β-trefoil core and a short C-terminus (Goldfarb, 2005) (Fig. 6*a*). Here we show that the long N-terminus of A-type FHFs, specifically residues 2-21, is the molecular component responsible for inducing TTX-resistant *I*_NaR_. While the F2A and F4A peptides show limited sequence similarity to the Navβ4 peptide, they all exhibit similar patterns of interactions with VGSCs. The F2A, F4A and Navβ4 peptides can all accelerate fast inactivation (likely through open-channel block), can be quickly expelled from the channel pore upon repolarization, and have positive and hydrophobic residues that seems to be essential for inducing *I*_NaR_ (Dover et al., 2010; Lewis and Raman, 2010; Venkatesan et al., 2014). These similarities lead us to propose that A-type FHFs induce *I*_NaR_ via the N-terminus by a relief-of-open-channel-block mechanism, similar to mechanism proposed for the Navβ4 peptide-dependent *I*_NaR_ (Lewis and Raman, 2014).

Previous studies proposed that FHF4 isoforms may regulate Nav1.6 *I*_NaR_ generation in Purkinje neurons (Yan et al., 2014; White et al., 2019). Although initially it was suggested that FHF4B indirectly enhanced *I*_NaR_ by attenuating inactivation of Nav1.6 (Yan et al., 2014), a more recent study indicated that FHF4A directly mediated Nav1.6 *I*_NaR_ ,showing a short peptide derived from a region of FHF4A adjacent to the β-trefoil core (residues 51-63) was able to induce robust *I*_NaR_ in FHF4A knockout mice (White et. Al, 2019). However, our data do not support the idea that FHF4A serves as a direct mediator of Nav1.6 *I*_NaR_, because FHF4A failed to mediate Nav1.6 *I*_NaR_ in our heterologous expression systems. Previously we demonstrated that FHF2A decreases Nav1.6 *I*_NaR_ in DRG neurons. In contrast, FHF2B, indirectly enhances *I*_NaR_ generation in DRG neurons (Barbosa et al., 2017). FHF2B lacks the long N-terminus of A-type FHFs but retains the β-trefoil core found in all FHFs that can bind to the cytoplasmic tail of VGSCs and destabilizes Nav1.6 fast-inactivation (also see Fig. 9h). Interestingly, FHF4 knockout accelerates the onset of fast inactivation of sodium currents (White et al., 2019) and hyperpolarizes the voltage-dependence of sodium current inactivation (Bosch et al., 2015) in cerebellar Purkinje neurons. This makes it possible that FHF4 isoforms regulate Nav1.6 *I*_NaR_ in neurons, at least in part, by reducing channel fast inactivation, instead of direct mediating *I*_NaR_. Computational modeling of *I*_NaR_ generation in mouse cerebellar Purkinje neurons suggests that blocking particle-independent mechanisms may account for TTX-sensitive *I*_NaR_ in some neuronal populations (Ransdell et al., 2022).

While neither full-length Navβ4 nor full-length FHF4A has been shown to induce Nav1.6 *I*_NaR_ in heterologous expression systems, our data show that FHF4A can induce long-term inactivation of Nav1.6 in ND7/23 cells (Fig. 10), which is consistent with previous studies demonstrating that A-type FHFs induce long-term inactivation of Nav1.6 and Nav1.5 (Dover et al., 2010; Yan et al., 2014; Yang et al., 2016; Barbosa et al., 2017). One possible explanation for the induction of long-term inactivation versus the induction of *I*_NaR_ generation is that A-type FHF N-terminus may bind more strongly to Nav1.6 and some other VGSCs than to Nav1.8 and Nav1.9. During repolarization, the driving force is only powerful enough to repel and unbind A-type FHF N-terminus from Nav1.8 and Nav1.9, but not from Nav1.6, and thus induces robust *I*_NaR_ from Nav1.8 and Nav1.9, but long-term inactivation in Nav1.6. However, we cannot rule out the possibility that post-translational modifications of either Nav1.6 or FHF4A allows FHF4A to directly induce *I*_NaR_ in Nav1.6, rather than inducing long-term inactivation, in specific neuronal populations such as cerebellar Purkinje neurons.

Intriguingly, our data show that heterologous expression of A-type FHF is sufficient to induce *I*_NaR_ in not only Nav1.8 and Nav1.9, but also, at least to some extent, in Nav1.5 and Nav1.7. This opens the door to further investigation of the molecular mechanism and the molecular manipulation of *I*_NaR_. Our data show that F4A peptide at up to 1 mM, a non-physiological concentration, is required to fully induce the same level of *I*_NaR_ as those induced by full-length FHF4A. This concentration is fivefold to tenfold higher than that of Navβ4 peptide (100 – 200 µM) that is required to reconstitute TTX-sensitive *I*_NaR_. The unusually high concentration apparently is due to the lack of the FHF β-trefoil core. The β-trefoil core does not directly generate *I*_NaR_, but it is likely crucial to facilitating *I*_NaR_ generation induced by N-terminus residues in A-type FHFs, as the core binding to the cytoplasmic tail of VGSCs (Liu et al., 2001; Goetz et al., 2009) would greatly raise the local concentration of the N-terminus near the channel pore. In addition, our data and that of others demonstrate that the β-trefoil domain shifts the voltage-dependence of steady-state inactivation in the positive direction, augmenting the “window currents” region (see Fig. 1*b*, *d*), where VGSCs activate but do not fully inactivate. Lewis and Raman (2013) showed that open-channel blockers might have higher affinity in VGSCs with DIVS4 deployed than with DIVS4 in the resting or partially deployed configuration. This also suggests that the molecular manipulation of *I*_NaR_ might be achieved by inhibiting the interaction of A-type FHF β-trefoil core with VGSC C-terminus.

Overall, our work substantially increases understanding of the role of A-type FHFs in sensory neuron excitability. We show that A-type FHFs exert various impact on neuronal excitability by differentially modulating the activities of VGSC isoforms. The accumulation of long-term inactivation seems to be the predominant effect of A-type FHFs on TTX-sensitive VGSC isoforms, although FHF-mediated *I*_NaR_ (only 0.3% of transient peak current) are inducible with some TTX-sensitive isoforms (Fig. 9*c,f*). By promoting long-term inactivation, FHF2A accumulatively decreases TTX-sensitive sodium currents (e.g. Nav1.6, Nav1.7) by > 20% (Venkatesan et al., 2014; Effraim et al., 2019). Prior studies demonstrated that A-type FHFs reduced action potential firing in hippocampal neurons, cerebellar granule neurons (Dover et al., 2010; Venkatesan et al., 2014) and in medium-sized DRG neurons, where Nav1.6 channels are predominantly expressed (Barbosa et al., 2017). Nav1.8 and Nav1.9 are two TTX-resistant subtypes mainly expressed in nociceptive sensory neurons (Fang et al., 2002; Cummins et al., 2007). In Fig. 6, we demonstrate that A-type FHF-mediated *I*_NaR_ significantly upregulate excitability of nociceptive DRG neurons. The *I*_NaR_ also result in broader action potentials and higher firing frequency. These observations are similar to those detected in Nav1.8 T790A-transfected DRG neurons, in which the T790A variant identified in the *Possum* transgenic mouse strain leads to increased *I*_NaR_ and enhanced excitability (Xiao et al., 2019). In addition to DRG neurons, Nav1.8/Nav1.9 have been colocalized with A-type FHFs within other neuronal populations, such as trigeminal ganglion neurons, myenteric neurons, magnocellular neurosecretory cells of the supraoptic nucleus, the outer layers of the substantia gelatinosa, and cerebellar neurons in animal models of multiple sclerosis (Craner et al., 2003; Vohra et al., 2006; Heanue et al., 2006; Huang et al., 2014; Osorio et al., 2014). This opens up the possibility that A-type FHF-mediated *I*_NaR_ extensively regulate excitability of the neurons throughout the PNS and CNS.

## Materials and Methods

### Plasmids, Sodium channel constructs and mutagenesis

Human FHF2A and FHF2B sequences were subcloned into pmTurquoise2-N1 vector as described by Barbosa et al. (2017). The pCMV6-AC-GFP plasmid encoding human FHF1A, FHF3A or FHF4A was purchased from Origene USA Technologies, Inc. (Rockville, MD). The cDNA construct encoding the human Nav1.5, mouse Nav1.8 and human Nav1.8 were subcloned into a pcDNA3.1 expression vector, respectively. The mutation N927K in Nav1.5 was constructed using the QuikChange XL (Stratagene) mutagenesis kit following the manufacturer’s instructions (Stratagene). Mutations were confirmed by sequencing. The scrambled shRNA and FHF4shRNA constructs were generously provided by Dr. Geoffrey S Pitt (Duke University). The scrambled shRNA and FHF4shRNA were subcloned into pAdTrack and pLVTHM vectors, respectively.

### Cell culture and transfection

Rat DRG neurons were acutely dissociated and cultured according to the procedure described previously (Xiao et al., 2019). Briefly, young adult (8 weeks) Sprague Dawley rats of either sex, in adherence with animal procedures approved by the Indiana University School of Medicine Institutional Animal Care and Use Committee, were killed by decapitation without anesthetization. All DRGs were removed quickly from the spinal cord and then incubated in DMEM containing collagenase (1 mg/ml) and protease (1 mg/ml). After the ganglia were triturated in DMEM supplemented with 10% FBS, cells were seeded on glass coverslips coated with poly-D-lysine and laminin. Cultures were maintained at 37°C in a 5% CO2 incubator. In order to be consistent with our previous studies (19), the Helios Gene Gun (Bio-Rad Laboratories) was used to transiently cotransfect rat DRG neurons. Cells were cotransfected with an internal ribosome entry site–EGFP (IRES-EGFP) vector plasmid (or an IRES-DsRed vector plasmid) containing a Nav1.8 shRNA targeting the rat Nav1.8 but not the codon optimized mouse Nav1.8 sequences. After transfection, DRG neurons were incubated in 10% FBS DMEM medium supplemented with mitotic inhibitors, 5-fluoro-2-deoxyuridine (50 µM, Sigma Aldrich) and uridine (150 µM, Sigma Aldrich), to prevent overgrowth of the supporting cells. DRG recordings were obtained from cells 2-5 days after transfection. Transfected cells were selected for recordings based on their ability to express EGFP. Under control conditions, the endogenous Nav1.8-type currents have an average current density of 947 ± 72 pA/pF (n = 70) and the Nav1.8 shRNA reduces endogenous Nav1.8-type current amplitudes in DRG neurons by 98% (Xiao et al., 2019).

Human Nav1.9, Nav1.9 K799N, Nav1.7 and Nav1.7 N945K channel cDNAs were stably expressed in the HEK-293-β1/β2 cell lines as described previously (Lin et al., 2016) and provided by Icagen Inc (Durham, North Carolina). They were incubated in 10% DMEM medium supplemented with G418 (400 mg/L) and puromycin (0.5 mg/L). Cell lines were transiently transfected by FHF1A, FHF2A, FHF2B, FHF3A or FHF4A using the Invitrogen Lipofectamine 2000. Nav1.9 cells were seeded on glass coverslips and incubated at 30 °C 24 hrs prior to patch-clamp recording.

HEK293 cells and ND7/23 cells were grown under standard tissue culture conditions (5% CO2 and 37 °C) in Dulbecco’s modified Eagle’s medium supplemented with 10% fetal bovine serum. Using the Invitrogen Lipofectamine 2000, human Nav1.5 and the mutant construct (N927K) were transiently co-transfected with FHF2B, FHF2A or FHF4A into HEK293 cells. The construct human Nav1.8 was transiently transfected into ND7/23 cells. The lipofectamine-DNA mixture was added to the cell culture medium and left for 3 h after which the cells were washed with fresh medium. Cells with green fluorescent protein fluorescence were selected for whole-cell patch clamp recordings 36-72 h after transfection.

### Electrophysiological recordings

Whole-cell voltage-clamp recordings were performed at room temperature (∼21 °C) using an EPC-10 amplifier and the Pulse program (HEKA Electronics). Recordings for hNav1.7 and hNav1.7 N945K were conducted at Icagen, Inc under similar conditions but with an Axopatch 200B amplifier and PCLAMP software (Molecular Devices).

For voltage clamp recordings, fire-polished electrodes (1.0 - 2.0 MΩ) were fabricated from 1.7 mm capillary glass using a P-97 puller (Sutter Instruments), and the tips were coated with sticky wax (KerrLab) to minimize capacitive artifacts and increase series resistance compensation. The pipette solution contained (in mM): 140 CsF, 1.1 EGTA, 10 NaCl and 10 HEPES, pH 7.3. The bathing solution was (in mM): 130 mM NaCl, 30 mM TEA Chloride, 1 mM MgCl_2_, 3 mM KCl, 1 mM CaCl_2_, 0.05 mM CdCl_2_, 10 mM HEPES, and 10 mM D-glucose, pH 7.3 (adjusted with NaOH). Tetrodotoxin (TTX; 500 nM) was added to the bath solution in order to block endogenous TTX-sensitive currents in DRG neurons, Nav1.9 and K799N stable cells, and cells expressing Nav1.8, Nav1.5 and the mutant N927K. The liquid junction potential for these solutions was < 8 mV; data were not corrected to account for this offset. The offset potential was zeroed before contacting the cell. After establishing the whole-cell recording configuration, the resting potential was held at -120 mV or -100 mV for 5 min to allow adequate equilibration between the micropipette solution and the cell interior. Linear leak subtraction, based on resistance estimates from four to five hyperpolarizing pulses applied before the depolarizing test potential, was used for all voltage clamp recordings. Membrane currents were usually filtered at 5 kHz and sampled at 20 kHz. Voltage errors were minimized using 70% - 90% series resistance compensation, and the capacitance artifact was canceled using the computer-controlled circuitry of the patch clamp amplifier.

*Steady-State activation*: Families of sodium currents were induced by 50-ms depolarizing steps to various potentials ranging from -120 to +40 mV in 5-mV (or 10-mV) increments. The conductance was calculated using the equation G(Nav) = I/(V - Vrev) in which I, V, and Vrev represent inward current value, membrane potential, and reversal potential, respectively.

*Steady-state inactivation*: The voltage dependence of steady-state inactivation was estimated using a standard double pulse protocol in which sodium currents were induced by a 50-ms depolarizing potential of 0 mV following a 500-ms prepulse at potentials that ranged from -130 to +10 mV with a 10-mV increment. Currents were plotted as a fraction of the maximum peak current. To obtain the midpoint voltages (V_1/2_) and slope factor (k), the curves of both steady-state activation and inactivation were fitted to a Boltzmann function.

*Recovery from inactivation*: Recovery from inactivation was assayed by the protocol that the cells were prepulsed to 0 mV for 50 ms to inactivate sodium channels and then brought back to -100 mV for increasing recovery durations before the test pulse to 0 mV.

*Resurgent currents*: *I*_NaR_ were elicited by repolarizing voltage steps from +10 mV to −100 for 100 ms (200 ms, or 1000 ms as indicated in Figures (*inset*)) in -5 mV increments, following a 20-ms depolarizing potential of +30 mV (or +100 mV). To avoid contamination from tail currents, Navβ4-induced Nav1.5 *I*_NaR_ were measured after 3.0 ms into the repolarization pulse, FHF-induced resurgent currents in Nav1.5, Nav1.7 and Nav1.9 were measured after 4.0 ms into the repolarization pulse, and FHF-induced Nav1.8 *I*_NaR_ were measured after 20 ms into the repolarization pulse. The relative *I*_NaR_ in Nav1.5, Nav1.7 and Nav1.8 were calculated by normalizing to the peak transient current elicited at 0 mV, but the relative of Nav1.9 resurgent currents were by normalizing to the peak transient current at -30 mV.

For current-clamp recordings, fire-polished electrodes (4.0 – 5.0 MΩ) were fabricated from 1.2 mm capillary glass using a P-97 (Sutter Instruments). The pipette solution contained the following (in mM): 140 KCl, 5 MgCl_2_, 5 EGTA, 2.5 CaCl_2_, 4 ATP, 0.3 GTP, and 10 HEPES, pH 7.3 (adjusted with KOH). The bathing solution contained the following (inmM): 140 NaCl, 1 MgCl2, 5 KCl, 2 CaCl_2_, 10 HEPES, and 10 glucose, pH 7.3 (adjusted with NaOH). Neurons were allowed to stabilize for 3 min in the current-clamp mode before initiating current injections to measure action potential activity.

### Immunocytochemistry

Immunocytochemistry was performed according the procedure as described previously (17). Briefly, the Helios Gene Gun (Bio-Rad Laboratories) was used to transiently transfect the scrambled shRNA, or FHF4shRNA in cultured DRG neurons. Three days after transfection, DRG neurons were fixed with 4% PFA (0.1 M phosphate buffer, pH 7.4) for 20 min and washed in PBS. Cells were then permeabilized in 1% Triton X-100 in PBS for 20 min at room temperature (∼21 °C), washed in PBS, blocked for 2 h (10% normal goat serum, 0.1% Triton X-100 in PBS) at room temperature, and washed with PBS. Cells were then incubated with monoclonal FHF4 antibody (1:200, N56/21, UC Davis/NIH NeuroMab Facility) or polyclonal anti-Navβ4 antibody (1:500, #Ab80539, Abcam) diluted in blocking solution over night at 4 °C. After additional PBS washes, cells were incubated with secondary antibody AlexaFluorTM Plus 555 Goat Anti-Mouse IgG (Invitrogen) in blocking solution at 1:1000 concentration for 2 h at room temperature. Coverslips were mounted in Prolong Gold Antifade (Invitrogen) and DRG neurons imaged using Leica Microscope system with a 20 objective (Biocompare). Images were analyzed with Leica software, and corrected mean cell fluorescence was calculated in Excel (Microsoft) by applying measurements obtained from image analysis using the equations: CMCF = (mean fluorescence intensity of selected cell) – (mean fluorescence of background).

### Experimental design and statistical analysis

The acquisition of control and experimental data was randomized. Data were analyzed using the software programs PulseFit (HEKA) and GraphPad Prism 5.0 (GraphPad Software, Inc., San Diego, CA). All data are shown as mean ± S.E. The number of separate experimental cells is presented as *n*. Statistical analysis was performed by Student’s t test, one-way ANOVA and χ^2^ analysis, and p < 0.05 indicated a significant difference.

## Acknowledgment

This work was supported, in whole or in part, by the National Institute of Neurological Disorders and Stroke of the National Institutes of Health under Award Numbers R21NS109896 (to Y. X. and T. R. C) and NS053422 (to T. R. C.), and the Indiana Spinal Cord & Brain Injury Research Fund from the Indiana State Department of Health (2020) (to Y. X.). We thank Dr. Geoffrey S. Pitt for generously providing the scrambled shRNA and FHF4shRNA constructs.

## Author Contributions

J.W.T and Y.P. performed experiments, analyzed data, and contributed to writing the manuscript, A.Z. and Z.L. performed experiments and analyzed data. Y.X. and T.R.C. conceived the study, oversaw the study, designed experiments and wrote the manuscript. Y.X. also performed experiments and analyzed data.

---

**Figure 2 - figure supplement 1.**
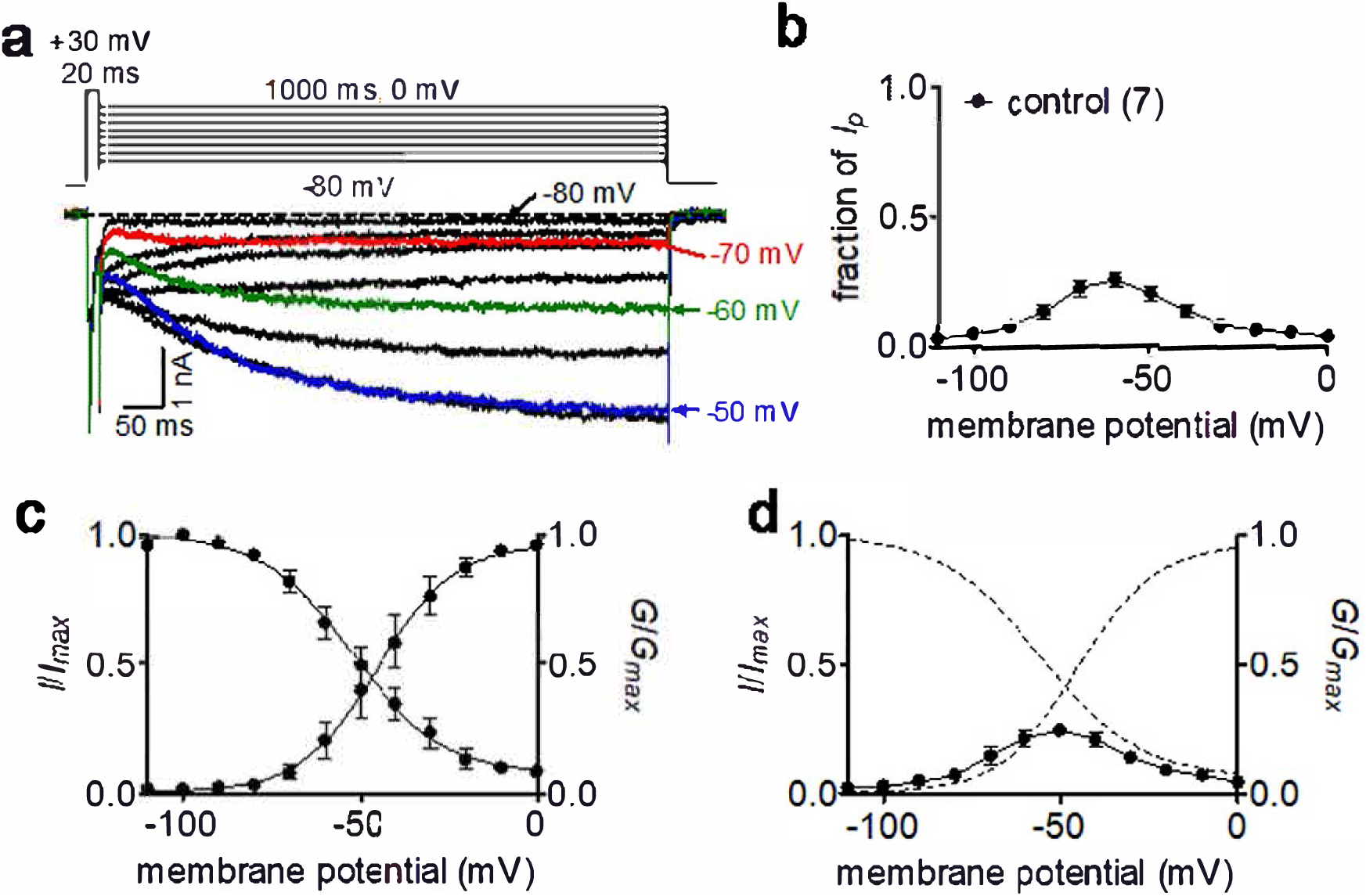
Extreme slow non-decay currents were caused by slow recovery from inactivation of Nav1.9 “window currents”. (**a**) Typical current traces elicited by a modified standard *I*_NaR_ protocol, in which repolarizing phase was extended to be 1000 ms (*inset*). (**b**) Normalization of the non-decay currents to the peak transient current with maximum amplitude. The non-decay currents were measured after 990 ms into the depolarizing pulse. (**c**) Normalized steady state activation and inactivation. (**d**) Overlay of the curves for normalized non-decay currents (filled circles), steady-state activation and inactivation (dash lines). The number of separate cells tested is indicated in parentheses.

**Figure 5 - figure supplement 1.**
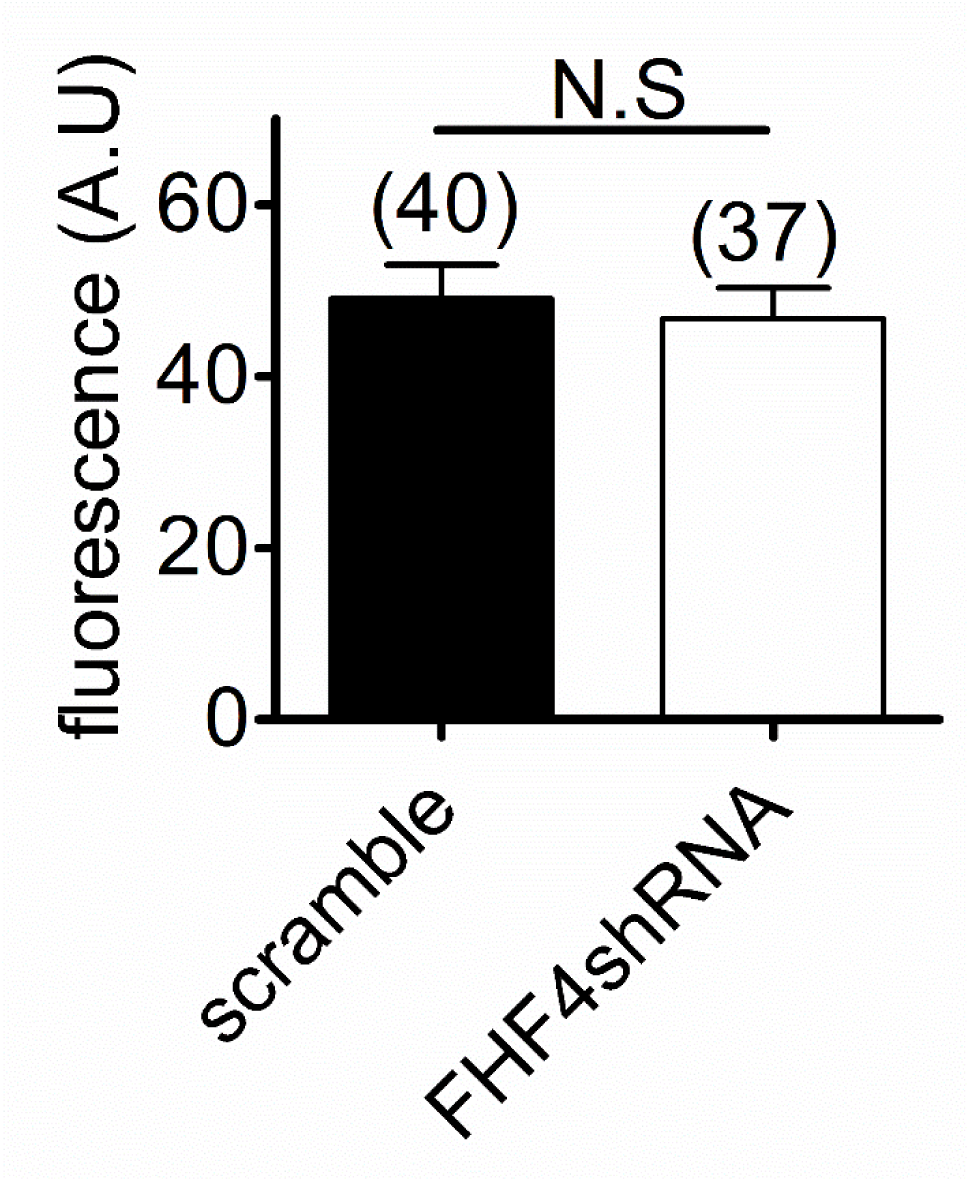
FHF4 knockdown did not influence Navβ4 expression in DRG neurons. The number of separate cells tested is indicated in parentheses. N.S, not significant.

**Figure 8 - figure supplement 1.**
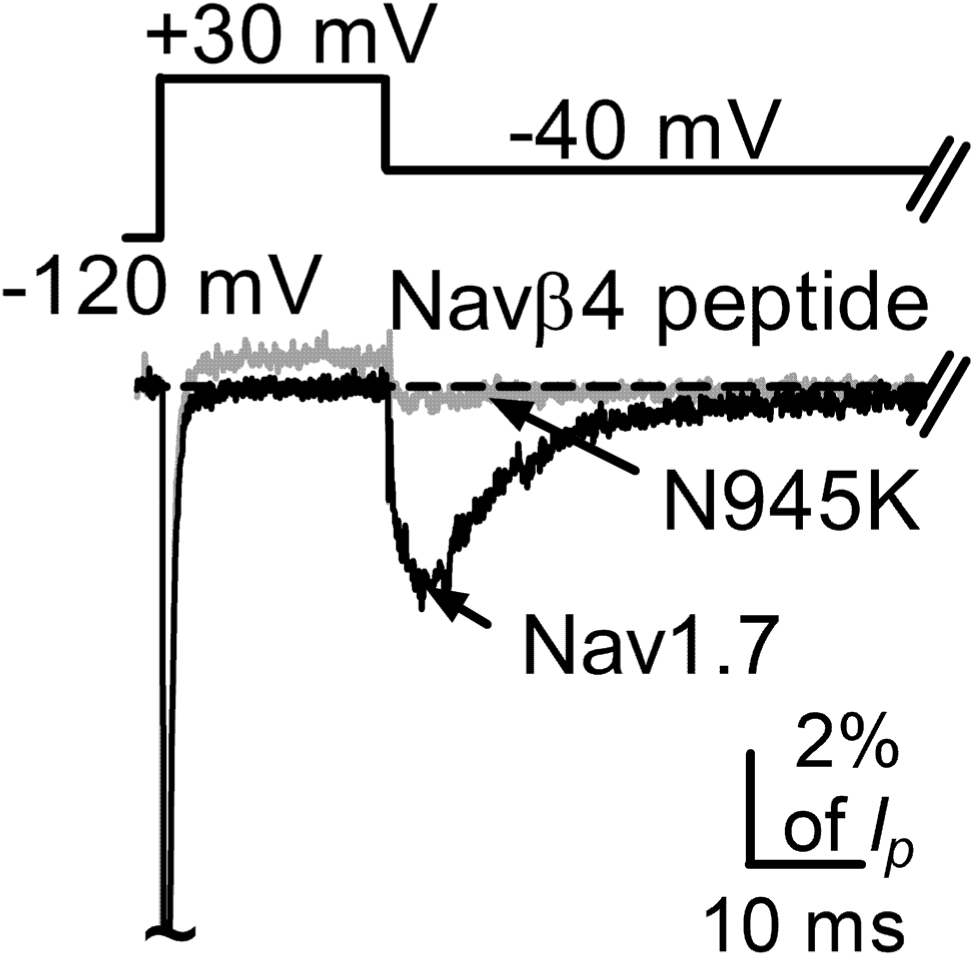
The N945K mutation substantially reduced Navβ4-mediated Nav1.7 *I*_NaR_ in HEK293 cells. Typical *I*_NaR_ traces in the presence of 200 µM Navβ4 peptide were elicited by the protocol shown in *inset*. Nav1.7, *black*; N945K, *grey*.

## References

1. Afshari FS, Ptak K, Khaliq ZM, Grieco TM, Slater NT, McCrimmon DR, Raman IM. 2004. Resurgent Na currents in four classes of neurons of the cerebellum. J. Neurophysiol., 92:2831–2843.

2. Aman TK, Grieco-Calub TM, Chen C, Rusconi R, Slat EA, Isom LL, Raman IM. 2009. Regulation of persistent Na current by interactions between beta subunits of voltage-gated Na channels. J. Neurosci., 29:2027–42.

3. Attwell D, Cohen I, Eisner D, Ohba M, Ojeda C. 1979. The steady state TTX-sensitive (“window”) sodium current in cardiac Purkinje fibres. Pflugers Arch, 379:137–142.

4. Bant JS, Raman IM. 2010. Control of transient, resurgent, and persistent current by open-channel block by Na channel beta4 in cultured cerebellar granule neurons. Proc Natl Acad Sci U S A. 107:12357–62.

5. Bant JS, Aman TK, Raman IM. 2013. Antagonism of lidocaine inhibition by open-channel blockers that generate resurgent Na current. J. Neurosci., 33:4976–87.

6. Barbosa C, Tan ZY, Wang R, Xie W, Strong JA, Patel RR, Vasko MR, Zhang JM, Cummins TR. 2015. Navβ4 regulates fast resurgent sodium currents and excitability in sensory neurons. Mol. Pain, 11:60.

7. Barbosa C, Xiao Y, Johnson AJ, Xie W, Strong JA, Zhang JM, Cummins TR. 2017. FHF2 isoforms differentially regulate Nav1.6-mediated resurgent sodium currents in dorsal root ganglion neurons. Pflugers Arch., 469:195–212.

8. Black JA, Dib-Hajj S, Baker D, Newcombe J, Cuzner ML, Waxman SG. 2000. Sensory neuron-specific sodium channel SNS is abnormally expressed in the brains of mice with experimental allergic encephalomyelitis and humans with multiple sclerosis. Proc. Natl. Acad. Sci. U S A, 97:11598–602.

9. Bosch MK, Carrasquillo Y, Ransdell JL, Kanakamedala A, Ornitz DM, Nerbonne JM. 2015. Intracellular FGF14 (iFGF14) Is Required for Spontaneous and Evoked Firing in Cerebellar Purkinje Neurons and for Motor Coordination and Balance. J Neurosci., 35:6752–6769.

10. Cannon SC, Bean BP. 2010. Sodium channels gone wild: resurgent current from neuronal and muscle channelopathies. J. Clin. Invest.,120:80–3.

11. Catterall WA, Goldin AL,, Waxman SG. 2005. International Union of Pharmacology. XLVII. Nomenclature and structure-function relationships of voltage-gated sodium channels. Pharmacol. Rev., 57:397–409.

12. Chen Y, Yu FH, Sharp EM, Beacham D, Scheuer T, Catterall WA. 2008. Functional properties and differential neuromodulation of Na(v)1.6 channels. Mol. Cell. Neurosci., 38:607–15.

13. Cox JJ, Reimann F, Nicholas AK, Thornton G, Roberts E, Springell K, Karbani G, Jafri H, Mannan J, Raashid Y, Al-Gazali L, Hamamy H, Valente EM, Gorman S, Williams R, McHale DP, Wood JN, Gribble FM, Woods CG. 2006. An SCN9A channelopathy causes congenital inability to experience pain. Nature, 444:894–8.

14. Craner MJ, Kataoka Y, Lo AC, Black JA, Baker D, Waxman SG. 2003. Temporal course of upregulation of Na(v)1.8 in Purkinje neurons parallels the progression of clinical deficit in experimental allergic encephalomyelitis. J. Neuropathol. Exp. Neurol., 62:968–975.

15. Cummins TR, Dib-Hajj SD, Waxman SG. 2004. Electrophysiological properties of mutant Nav1.7 sodium channels in a painful inherited neuropathy. J. Neurosci., 24:8232–8236.

16. Cummins TR, Sheets PL, Waxman SG. 2007. The roles of sodium channels in nociception: Implications for mechanisms of pain. Pain, 131:243–57.

17. Dib-Hajj SD, Cummins TR, Black JA, Waxman SG. 2010. Sodium channels in normal and pathological pain. Annual Review of Neuroscience, 33:325–347.

18. Dib-Hajj SD, Black JA, Waxman SG. 2015. NaV1.9: a sodium channel linked to human pain. Nat. Rev. Neurosci., 16:511–9.

19. Dover K, Solinas S, D’Angelo E, Goldfarb M. 2010. Long-term inactivation particle for voltage-gated sodium channels. J. Physiol., 588:3695–711.

20. Effraim PR, Huang J, Lampert A, Stamboulian S, Zhao P, Black JA, Dib-Hajj SD, Waxman SG. 2019. Fibroblast growth factor homologous factor 2 (FGF-13) associates with Nav1.7 in DRG neurons and alters its current properties in an isoform-dependent manner. Neurobiol Pain, 6:100029.

21. Enomoto A, Han JM, Hsiao CF, Wu N, Chandler SH. 2006.Participation of sodium currents in burst generation and control of membrane excitability in mesencephalic trigeminal neurons. J. Neurosci., 26:3412–3422.

22. Fang X, Djouhri L, Black JA, Dib-Hajj SD, Waxman SG, Lawson SN. 2002. The presence and role of the tetrodotoxin-resistant sodium channel Na(v)1.9 (NaN) in nociceptive primary afferent neurons. J Neurosci, 22:7425–7433.

23. Goetz R, Dover K, Laezza F, Shtraizent N, Huang X, Tchetchik D, Eliseenkova AV, Xu CF, Neubert TA, Ornitz DM, Goldfarb M, Mohammadi M. 2009. Crystal structure of a fibroblast growth factor homologous factor (FHF) defines a conserved surface on FHFs for binding and modulation of voltage-gated sodium channels. J. Biol. Chem., 284:17883–96.

24. Goldfarb M. 2005. Fibroblast growth factor homologous factors: evolution, structure, and function. Cytokine Growth Factor Rev., 16:215–20.

25. Grieco TM, Raman IM. 2004. Production of resurgent current in NaV1.6-null Purkinje neurons by slowing sodium channel inactivation with beta-pompilidotoxin. J Neurosci. 24:35–42.

26. Grieco TM, Malhotra JD, Chen C, Isom LL, Raman IM. 2005. Open-channel block by the cytoplasmic tail of sodium channel beta4 as a mechanism for resurgent sodium current. Neuron, 45: 233–44.

27. Heanue TA, Pachnis V. 2006. Expression profiling the developing mammalian enteric nervous system identifies marker and candidate Hirschsprung disease genes. Proc. Natl. Acad. Sci. U S A, 103:6919–24.

28. Huang J, Yang Y, Zhao P, Gerrits MM, Hoeijmakers JG, Bekelaar K, Merkies IS, Faber CG, Dib-Hajj SD, Waxman SG. 2013. Small-fiber neuropathy Nav1.8 mutation shifts activation to hyperpolarized potentials and increases excitability of dorsal root ganglion neurons. J. Neurosci., 33:14087–97.

29. Huang J, Han C, Estacion M, Vasylyev D, Hoeijmakers JG, Gerrits MM, Tyrrell L, Lauria G, Faber CG, Dib-Hajj SD, Merkies IS, Waxman SG, PROPANE Study Group. 2014. Gain-of-function mutations in sodium channel Na(v)1.9 in painful neuropathy. Brain, 137:1627–42.

30. Jarecki BW, Piekarz AD, Jackson JO, Cummins TR. 2010. Human voltage-gated sodium channel mutations that cause inherited neuronal and muscle channelopathies increase resurgent sodium currents. J Clin Invest, 120:369–78.

31. Kim JH, Kushmerick C, von Gersdorff H. 2010. Presynaptic resurgent Na+ currents sculpt the action potential waveform and increase firing reliability at a CNS nerve terminal. J. Neurosci., 30:15479–15490.

32. Lewis AH, Raman IM. 2011. Cross-species conservation of open-channel block by Na channel β4 peptides reveals structural features required for resurgent Na current. J. Neurosci., 31:11527–36.

33. Lewis AH, Raman IM. 2013. Interactions among DIV voltage-sensor movement, fast inactivation, and resurgent Na current induced by the NaV4 open-channel blocking peptide. J. Gen. Physiol., 142:191–206.

34. Lewis AH, Raman IM. 2014. Resurgent current of voltage-gated Na(+) channels. J. Physiol., 592, 4825–38.

35. Li GD, Wo Y, Zhong MF, Zhang FX, Bao L, Lu YJ, Huang YD, Xiao HS, Zhang X. 2002. Expression of fibroblast growth factors in rat dorsal root ganglion neurons and regulation after peripheral nerve injury. Neuroreport, 13:1903–7.

36. Lin Z, Santos S, Padilla K, Printzenhoff D, Castle NA. 2016. Biophysical and Pharmacological Characterization of Nav1.9 Voltage Dependent Sodium Channels Stably Expressed in HEK-293 Cells. PLoS One, 11:e0161450.

37. Liu CJ, Dib-Hajj SD, Waxman SG. 2001. Fibroblast growth factor homologous factor 1B binds to the C terminus of the tetrodotoxin-resistant sodium channel rNav1.9a (NaN). J. Biol. Chem., 276:18925–33.

38. Lou JY, Laezza F, Gerber BR, Xiao M, Yamada KA, Hartmann H, Craig AM, Nerbonne JM, Ornitz DM. 2005. Fibroblast growth factor 14 is an intracellular modulator of voltage-gated sodium channels. J. Physiol., 569:179–93.

39. Miyazaki H, Oyama F, Inoue R, Aosaki T, Abe T, Kiyonari H, Kino Y, Kurosawa M, Shimizu J, Ogiwara I, Yamakawa K, Koshimizu Y, Fujiyama F, Kaneko T, Shimizu H, Nagatomo K, Yamada K, Shimogori T, Hattori N, Miura M, Nukina N. (2014) Singular localization of sodium channel β4 subunit in unmyelinated fibres and its role in the striatum. Nat Commun. 21;5:5525.

40. Namadurai S, Yereddi NR, Cusdin FS, Huang CL, Chirgadze DY, Jackson AP. 2015. A new look at sodium channel β subunits. Open Biol., 5:140–192.

41. Osorio N, Korogod S, Delmas P. 2014. Specialized functions of Nav1.5 and Nav1.9 channels in electrogenesis of myenteric neurons in intact mouse ganglia. J. Neurosci., 34:5233–44.

42. Pan Y, Cummins TR. 2020. Distinct functional alterations in SCN8A epilepsy mutant channels. J Physiol. 598:381–401.

43. Patel RR, Barbosa C, Brustovetsky T, Brustovetsky N, Cummins TR. 2016 Aberrant epilepsy-associated mutant Nav1.6 sodium channel activity can be targeted with cannabidiol. Brain, 139:2164–81.

44. Raman IM, Bean BP. 1997. Resurgent sodium current and action potential formation in dissociated cerebellar Purkinje neurons. J. Neurosci., 17:4517–4526.

45. Ransdell JL, Dranoff E, Lau B, Lo WL, Donermeyer DL, Allen PM, Nerbonne JM. 2017. Loss of Navβ4-Mediated Regulation of Sodium Currents in Adult Purkinje Neurons Disrupts Firing and Impairs Motor Coordination and Balance. Cell Rep., 20:1502.

46. Ransdell JL, Moreno JD, Bhagavan D, Silva JR, Nerbonne JM. 2022. Intrinsic mechanisms in the gating of resurgent Na^+^ currents. Elife 11:e70173.

47. Rush AM, Wittmack EK, Tyrrell L, Black JA, Dib-Hajj SD, Waxman SG. 2006. Differential modulation of sodium channel Na(v)1.6 by two members of the fibroblast growth factor homologous factor 2 subfamily. Eur J Neurosci., 23:2551–2562.

48. Tanaka BS, Zhao P, Dib-Hajj FB, Morisset V, Tate S, Waxman SG, Dib-Hajj SD. 2016. A gain-of-function mutation in Nav1.6 in a case of trigeminal neuralgia. Mol. Med., 22.

49. Theile JW, Jarecki BW, Piekarz AD, Cummins TR. 2011. Nav1.7 mutations associated with paroxysmal extreme pain disorder, but not erythromelalgia, enhance Navbeta4 peptide-mediated resurgent sodium currents. J. Physiol., 589:597–608.

50. Venkatesan K, Liu Y, Goldfarb M. 2014. Fast-onset long-term open-state block of sodium channels by A-type FHFs mediates classical spike accommodation in hippocampal pyramidal neurons. J. Neurosci., 34:16126–39.

51. Vohra BP, Tsuji K, Nagashimada M, Uesaka T, Wind D, Fu M, Armon J, Enomoto H, Heuckeroth RO. 2006. Differential gene expression and functional analysis implicate novel mechanisms in enteric nervous system precursor migration and neuritogenesis. Dev Biol, 298:259–71.

52. White HV, Brown ST, Bozza TC, Raman IM. 2019. Effects of FGF14 and Na(V)β4 deletion on transient and resurgent Na current in cerebellar Purkinje neurons. J. Gen. Physiol., 151:1300–1318.

53. Wildburger NC, Ali SR, Hsu WC, Shavkunov AS, Nenov MN, Lichti CF, LeDuc RD, Mostovenko E, Panova-Elektronova NI, Emmett MR, Nilsson CL, Laezza F. 2015. Quantitative proteomics reveals protein-protein interactions with fibroblast growth factor 12 as a component of the voltage-gated sodium channel 1.2 (nav1.2) macromolecular complex in Mammalian brain. Mol. Cell Proteomics, 14:1288–300.

54. Wittmack EK, Rush AM, Craner MJ, Goldfarb M, Waxman SG, Dib-Hajj SD. 2004. Fibroblast growth factor homologous factor 2B: association with Nav1.6 and selective colocalization at nodes of Ranvier of dorsal root axons. J. Neurosci., 24:6765–75.

55. Wang C, Hennessey JA, Kirkton RD, Wang C, Graham V, Puranam RS, Rosenberg PB, Bursac N, Pitt GS. 2011a. Fibroblast growth factor homologous factor 13 regulates Na+ channels and conduction velocity in murine hearts. Circ. Res., 109:775–82.

56. Wang C, Wang C, Hoch EG, Pitt GS. 2011b. Identification of novel interaction sites that determine specificity between fibroblast growth factor homologous factors and voltage-gated sodium channels. J. Biol. Chem., 286:24253–63.

57. Xiao Y, Barbosa C, Pei Z, Xie W, Strong JA, Zhang JM, Cummins TR. 2019. Increased Resurgent Sodium Currents in Nav1.8 Contribute to Nociceptive Sensory Neuron Hyperexcitability Associated with Peripheral Neuropathies. J. Neurosci., 39:1539–1550.

58. Xie W, Tan ZY, Barbosa C, Strong JA, Cummins TR, Zhang JM. 2016. Upregulation of the sodium channel NaVbeta4 subunit and its contributions to mechanical hypersensitivity and neuronal hyperexcitability in a rat model of radicular pain induced by local dorsal root ganglion inflammation. Pain, 157:879–891.

59. Yan H, Pablo JL, Wang C, Pitt GS. 2014. FGF14 modulates resurgent sodium current in mouse cerebellar Purkinje neurons. Elife, 3, e04193.

60. Yang J, Wang Z, Sinden DS, Wang X, Shan B, Yu X, Zhang H, Pitt GS, Wang C. 2016. FGF13 modulates the gating properties of the cardiac sodium channel Na(v)1.5 in an isoform-specific manner. Channels (Austin), 10:410–420.

